# Mechanistic dissection of GRHL2 and PR transcriptional co-regulation in breast cells

**DOI:** 10.1101/2025.09.01.673508

**Authors:** M.T. Aarts, A. Nordin, C. Cantù, A.L. van Boxtel, R. van Amerongen

## Abstract

Gene expression is controlled by complex transcriptional networks in which transcription factors and their cognate enhancer elements integrate developmental and environmental cues. The progesterone receptor (PR), a hormone-activated transcription factor, is essential for breast development and physiology, yet how it engages with the chromatin and lineage-specific cofactors remains unclear. Using an unbiased approach, we identify the epithelial transcription factor grainyhead-like 2 (GRHL2) as a key co-regulator of PR activity in hormone responsive breast cancer cells. We show that GRHL2 interacts with PR in a progesterone-independent manner. Upon progesterone stimulation, GRHL2 and PR are both recruited to distal enhancer elements of target genes. Furthermore, GRHL2- and PR-bound elements connect spatially through chromatin looping to regulate shared targets. These findings uncover a previously unrecognized mechanism by which GRHL2 and PR coordinate gene regulation through both chromatin binding and 3D genome architecture modification, positioning GRHL2 as a crucial modulator of steroid hormone receptor function.

## Introduction

Gene expression is tightly regulated by complex transcriptional networks, in which DNA-binding transcription factors (TFs) play a central role. These networks govern when and where genes are expressed by integrating cell-intrinsic developmental cues and extrinsic environmental signals^1^. TFs bind to distinct motifs in regulatory DNA elements to modulate transcriptional output^2–4^. Rather than acting alone, TFs operate in combinatorial assemblies with additional transcription factors, co-factors and chromatin remodelers. This layered regulatory architecture ensures that distinct gene expression programs are executed in a cell type– and context-specific manner, enabling the emergence of specialized cellular identities throughout development and their faithful maintenance in adult tissues^5,6^.

Steroid hormone receptors represent a unique class of transcription factors, distinguished by their dual role as ligand-activated receptors and direct modulators of gene expression. Unlike many transcription factors with constitutive DNA binding activity, steroid receptor function is extrinsically regulated by the presence of specific hormones. Upon ligand binding, these receptors accumulate in the nucleus, where they dynamically associate with hormone response elements at the chromatin to regulate transcriptional programs^7^.

This class of receptors includes the estrogen receptor (ER), which has its own unique DNA binding motif, and the 3-keto-steroid receptors, which share a consensus sequence: progesterone receptor (PR), mineralocorticoid receptor (MR), glucocorticoid receptor (GR) and androgen receptor (AR). Activated by ovarian hormones, ER and PR are key regulators of postnatal breast tissue development^8–10^. In this tissue, they are almost exclusively expressed in a subset of luminal cells, named the mature luminal or hormone-sensing cells. The majority of attention has been directed to resolving the signaling mechanism of ER, given its role (and targetability) in hormone-dependent breast cancer. The molecular mechanisms of PR signaling remain less well understood, but it regulates multiple cell intrinsic and paracrine signaling pathways. These pathways promote local proliferation and basement membrane restructuring^11^ which in turn support ductal outgrowth, side branching and alveologenesis during pregnancy^10^. As such, PR is crucial for breast development and physiology and antiprogestins are actively explored from the perspective of breast cancer prevention and treatment^12^.

Steroid receptors such as ER and PR do not act in isolation but function within complex regulatory networks involving additional associated proteins. Among these, pioneer factors, a class of transcription factors capable of modulating chromatin accessibility^13^, have increasingly been recognized as critical co-regulators of steroid receptor activity, particularly in hormone-dependent cancers^14^. For instance, ER cooperates with pioneer factors such as FOXA1 and GATA3 to facilitate transcriptional activation of its target genes^15–20^. In contrast, the extent to which PR engages in similar interactions remains poorly understood. Grainyhead-like 2 (GRHL2) has recently emerged as a potential pioneer factor in hormone receptor-positive cancers, including breast cancer^21^. However, nearly all studies to date have focused on GRHL2 in the context of ER and estrogen signaling, leaving its role in PR- and progesterone-mediated regulation unexplored^22–26^.

In this study, we identify and dissect the functional transcriptional interaction of PR and GRHL2, a member of the GRHL protein family. This family includes three evolutionarily conserved pioneer transcription factors (GRHL1, GRHL2, and GRHL3) with important roles in epithelia^27,28^. They share a highly conserved DNA-binding domain and display substantial structural homology^29,30^. Despite these similarities, the GRHL family members exhibit distinct spatiotemporal expression patterns and execute unique epithelial transcriptional programs that underpin their specific biological functions.

Here we combine genomic, transcriptomic and proteomic, approaches to provide the first mechanistic dissection of the coordinate transcriptional activities of PR and GRHL2 in human breast cancer cells. We show that PR and GRHL2 can interact, independently of progesterone, and frequently co-occupy enhancer elements upon progesterone stimulation. Moreover, GRHL2 is required for the regulation of a set of PR target genes. Chromatin looping analysis further revealed how both shared (i.e. to the same DNA element) and distinct (i.e. to linearly separated DNA elements) PR and GRHL2 binding events can converge over long distances in 3D chromatin space to regulate common targets. Together, our findings uncover a new layer of PR gene regulation, shaped by GRHL2 and the 3D genome architecture.

## Results

### GRHL2 physically interacts with PR in a progesterone independent manner

To identify nuclear interactors of GRHL2 in an unbiased manned, we performed Rapid immunoprecipitation mass spectrometry of endogenous protein (RIME)^31^ in T47DS cells (**Fig1A**). Under stripped-serum, non-hormone stimulated conditions we identified 1,140 nuclear GRHL2-associated proteins, including several previously reported interactors such as FOXA1^18^, KMT2C/D (MLL3/4)^32^, GRHL1^27,33^ (**Fig1B**). Of these 1,140 interactors, a total of 103 were annotated as transcription factors or chromatin-associated/modifying proteins, based on the Panther Classification System^34^. We ranked these proteins by protein score to identify and highlight high-confidence candidates (**Fig1C**). Interestingly, several of the top-scoring interactors, such as CTBP1, MTA2, SMARCC2, SMARCD2, CHD3, and PHF6, are involved in chromatin accessibility and remodeling, supporting the role of GRHL2 as a pioneering factor and modulator of chromatin accessibility^28^ (**Fig1C**).

**Fig 1.**
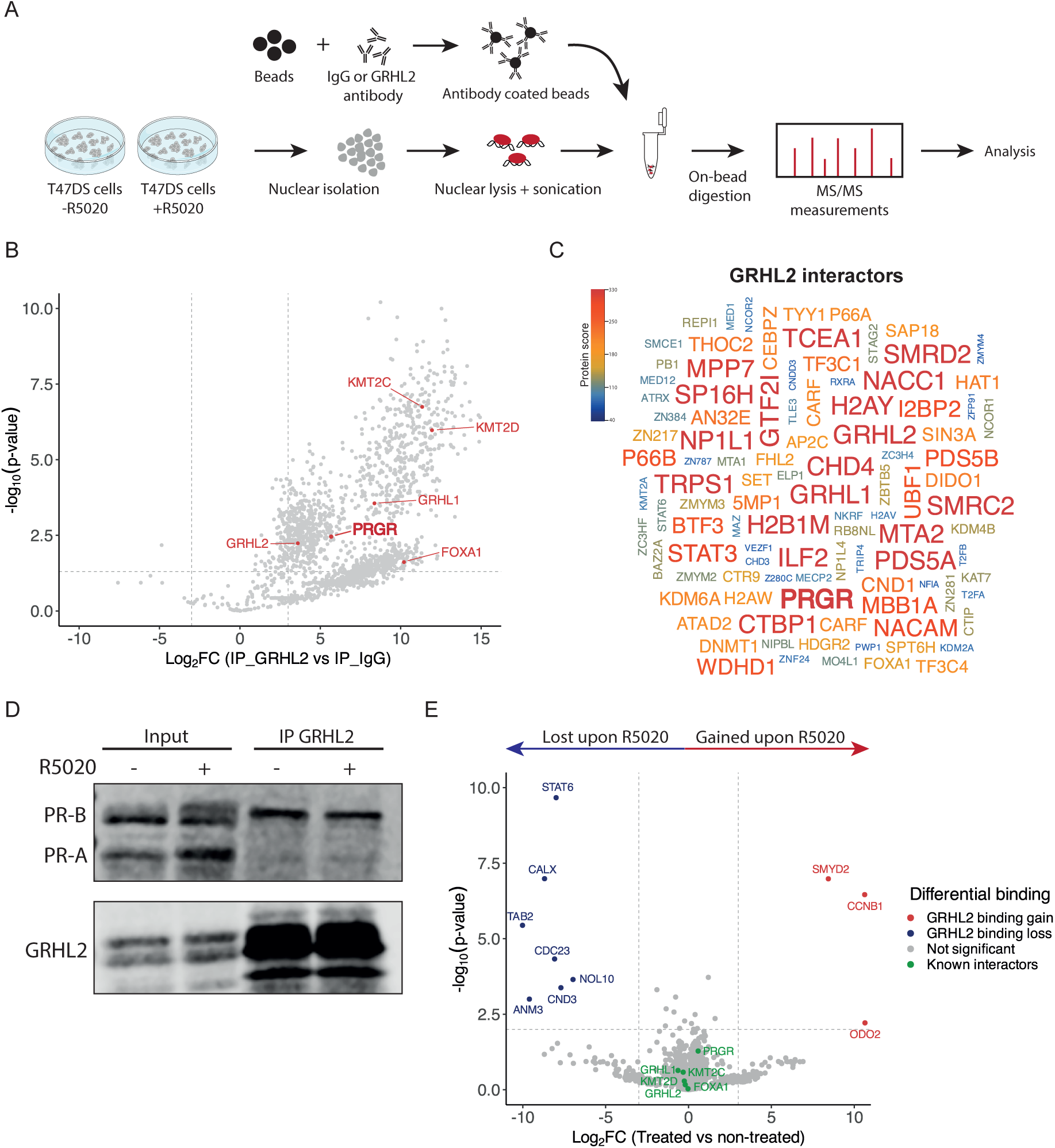
RIME analysis of GRHL2-associated proteins reveals GRHL2-PR interaction independent of progesterone. A) Schematic overview of the Rapid immunoprecipitation mass spectrometry of endogenous protein (RIME)^31^ protocol used to identify GRHL2 interactors in hormone depleted and progesterone treated conditions. B) Volcano plot depicting the results of the GRHL2 RIME of hormone depleted conditions vs. the IgG control. Each grey dot represents a single protein identified in mass spectrometry. Grey dotted lines represent Log2FC (−3,3) and -Log10(p-value) cutoffs (0.05). A total of 1,140 proteins passed these criteria and were thus identified as a GRHL2 interactor. Highlighted dots in red are a few examples of previously reported GRHL2 interactors^18,27,32,33,35^. C) Word cloud presenting 103 GRHL2 interactors filtered as transcription factors or chromatin associated/modifying proteins according to the Panther Classification System^34^. The size and the color of the protein names represent the confidence of the identified interaction based on the protein score. D) Western blot of co-immunoprecipitation validation of the GRHL2 - PR interaction in hormone depleted and 4 hours 1 nM of the PR agonist R5020 stimulated conditions. Western blot shows GRHL2 and PR, isoform A and B, protein of input and immunoprecipitated samples. E) Volcano plot depicting the results of the GRHL2 RIME in hormone depleted vs. 4 hour 1 nM R5020 treated conditions. Each grey dot represents a single protein identified in mass spectrometry. Grey dotted lines represent Log2FC (- 3,3) and -Log10(p-value) cutoffs (0.01). Only 10 proteins (7 lost, 3 gained) passed these criteria and were identified as a progesterone dependent GRHL2 interactor. Highlighted dots in blue are proteins that lost GRHL2, highlighted dots in red are proteins that gained GRHL2 binding.

Unexpectedly, PR emerged as one of the most robust GRHL2 interactors in these stripped-serum conditions, suggesting a hormone-independent association between the two factors (**Fig1B-C**). We validated this interaction by co-immunoprecipitation in both hormone-stripped vehicle-treated and hormone-stripped R5020-treated T47DS cells, confirming that the GRHL2–PR interaction remained unaffected by progesterone stimulation (**Fig1D**). Interestingly, we find that GRHL2 specifically binds to the full-length and transcriptionally active isoform of PR, PR-B, but not to PR-A (**Fig1D**).

Since the GRHL2-PR interaction is not dependent on the presence of progesterone, we wondered if and how progesterone altered the GRHL2 interactome. Therefore, we also performed the GRHL2 RIME in T47DS cells following 4 hour treatment with the synthetic progestin R5020 (**Fig1E**). Comparison with the untreated condition revealed that only few interactions were affected. For seven proteins, the interaction with GRHL2 was lost upon progesterone stimulation. Conversely, three proteins were recruited only in the presence of hormone (**Fig1E**). With the exception of STAT6 (interaction loss) and SMYD2 (interaction gain), these differentially interacting proteins were not classified as transcription factors or chromatin regulators, and no direct relation to progesterone or PR has thus far been reported in the literature. Since the GRHL2-PR interaction itself has also not been functionally characterized, we focused our subsequent analyses on this potential transcriptional partnership.

### Global mapping of GRHL2 and PR DNA binding reveals co-occupancy at enhancer sites

To determine whether the GRHL2–PR interaction occurs at shared genomic loci, we re-analyzed two previously published ChIP-seq datasets for GRHL2^23^ and PR^36^ in T47D cells. Of note, the GRHL2 dataset was generated under stripped-serum conditions, thus representing GRHL2 binding under estrogen and progesterone depleted conditions. The PR dataset was also generated under stripped-serum conditions, but here the cells were stimulated with 1 nM R5020. In this analysis, we identified 46,746 PR and 28,925 GRHL2 peaks. Of these, 6,335 sites were overlapping, representing 13.5% of the PR and 21.9% of the total GRHL2 binding events (**Fig2A**). One example of such a GRHL2-PR shared region is shown (**Fig2B**). Heatmap visualization of PR and GRHL2 binding at PR-only, GRHL2-only and GRHL2-PR overlapping sites revealed that the strongest PR and GRHL2 signal is concentrated at overlapping regions, although some GRHL2 signal is also detected at the strongest PR peaks and vice versa (**Fig2C-D**). The observation that GRHL2 and PR signals co-occur beyond the defined overlapping peaks suggests that their functional cooperation likely extends beyond the stringently thresholded sites identified in our analysis (**Fig2C-D**).

**Fig 2.**
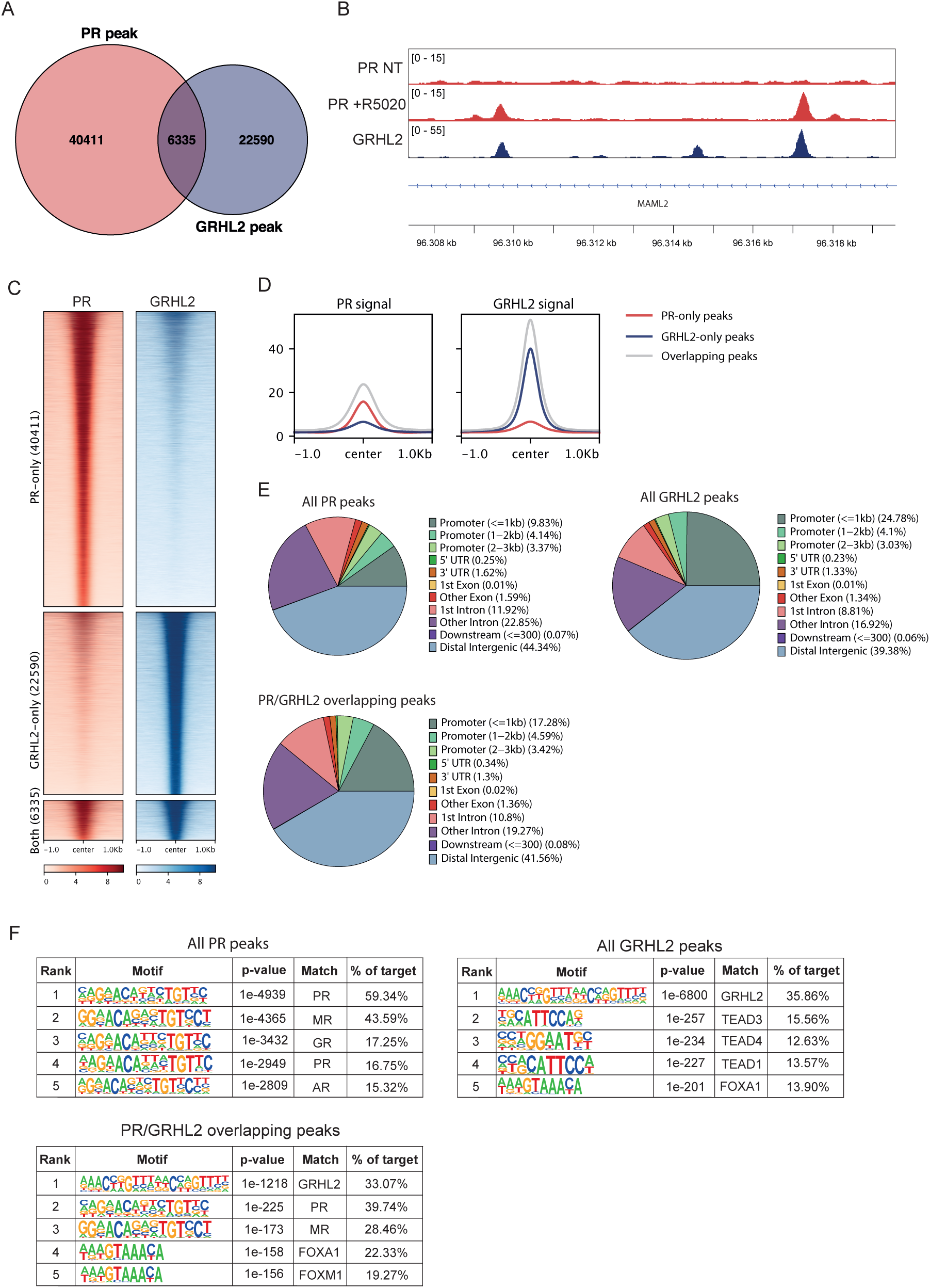
GRHL2 and PR co-bind regulatory regions across the genome. A) Venn diagram showing the overlap between GRHL2^23^ and 1 nM R5020 stimulated PR^36^ ChIP-seq peaks. Peaks were considered overlapping if they overlapped by at least 1 bp. B) A representative example of two PR and GRHL2 overlapping peaks. Data was visualized in the IGV browser^37^. NT= non-treated. C) Heatmaps showing binding ChIP-seq signal of GRHL2^23^ and 1 nM R5020 stimulated PR^36^ at PR-only, GRHL2-only and GRHL2-PR co-occupied genomic regions. D) Signal intensity plots showing the cumulative ChIP-seq signal of GRHL2^23^ and 1 nM R5020 stimulated PR^36^ at the PR-only, GRHL2-only and GRHL2-PR co-occupied genomic sites. E) Genomic annotation of all PR, all GRHL2 or GRHL2-PR shared peaks by ChIPseeker^38,39^. Sites were annotated to 5’UTR, promotors, 1st exon, other exons, 1st intron, other introns, 3’UTR introns, distal intergenic regions and downstream. Promotor regions were divided into three subcategories defines as <= 1 kb from the TSS, 1-2 kb from the TSS or 2-3 kb from the TSS. F) Top five transcription factor motifs that were determined as enriched in all 1 nM R5020 stimulated PR peaks, all GRHL2 peaks or the GRHL2-PR shared peaks as determined by HOMER motif analysis^40^.

Genomic annotation of the PR and GRHL2 ChIP-seq peaks showed that the majority of the PR (∼66%) and GRHL2 (∼57%) sites, including the overlapping PR-GRHL2 (∼61%) sites are located at enhancers in the distal intergenic or intronic regions (**Fig2E**). Notably, GRHL2 peaks were more likely located within 1 kb of a transcription start site (TSS) than PR containing peaks (∼25% of all GRHL2 peaks and ∼17% of GRHL2-PR shared sites, compared to <10% of all PR peaks) (**Fig2E**). Motif analysis confirmed strong enrichment for PR and GRHL2 motifs in both their individual and overlapping peaks (**Fig2F**). Moreover, PR or GRHL2 bound, as well as GRHL2-PR overlapping regions were enriched for FOXA1 motifs. In addition, TEAD motifs were specifically enriched in GRHL2 bound, including GRHL2-PR overlapping sites. This suggests that both FOXA1 and TEAD act on similar regulatory regions as GRHL2 and PR in breast tissue (**Fig2F**).

We next asked if and how progesterone would modulate chromatin binding of GRHL2. To obtain more insight in such binding dynamics, we performed CUT&RUN analysis for GRHL2 and PR in hormone deprived T47DS cells under both unstimulated and progesterone-stimulated conditions. Without hormone stimulation, 7,039 GRHL2 peaks were identified, whereas 5,520 GRHL2 peaks were identified following 4 hours of R5020 treatment (**Fig3A**). Only 713 of these sites overlapped. Moreover, the GRHL2 signal significantly increases upon R5020 stimulation, indicating that GRHL2 chromatin binding is redistributed and strengthened upon R5020 stimulation (**Fig3A**). This effect is unlikely to be fully explained by a mere increase in GRHL2 protein levels, as 4 hour R5020 treatment elevated GRHL2 protein levels only slightly, with at most a 20% (and statistically insignificant) increase in protein abundance (**SupFig2A-B**). For PR, we identified 3,033 peaks after 4 hours of R5020 stimulation. Overlapping the GRHL2 and PR peaks revealed that GRHL2 and PR binding sites only overlap under R5020-treated condition (158 sites, **Fig3B**, **SupFig1A-B**). Motif analysis of these sites confirmed enrichment for both PR and GRHL2 motifs, as expected (**Fig3C**). Given that PR and GRHL2 interact in both the absence and presence of progesterone (**Fig1**), this suggests that PR activation facilitates GRHL2 recruitment to the progesterone responsive chromatin elements.

**Fig 3.**
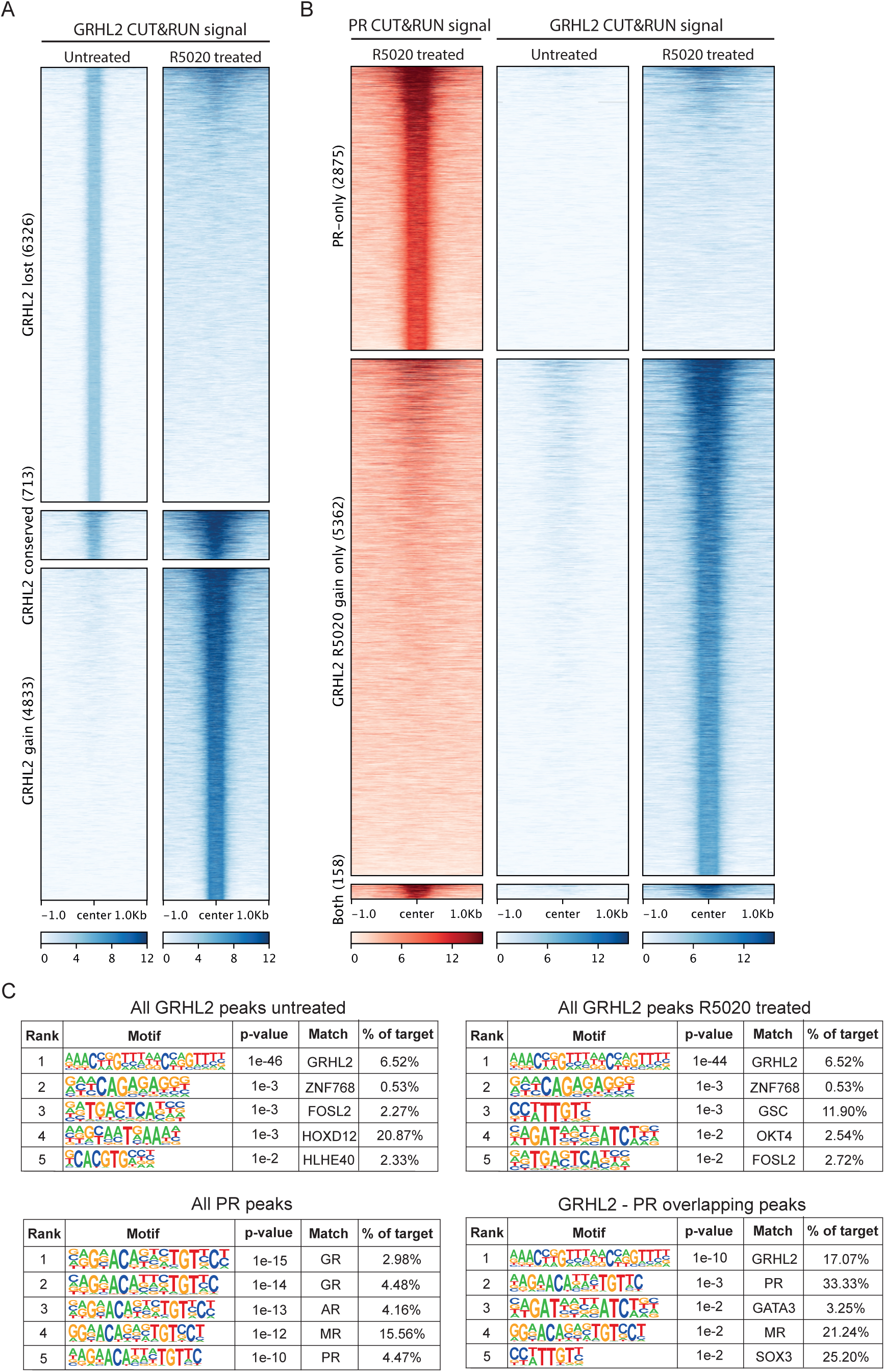
CUT&RUN reveals GRHL2 redistribution upon progesterone stimulation. A) Heatmaps showing CUT&RUN GRHL2 signal in untreated and 4 hour 1 nM R5020 stimulated conditions at sites identified as GRHL2 bound in hormone depleted (GRHL2 lost), GRHL2 bound in hormone depleted and R5020 stimulated conditions (GRHL2 conserved), or GRHL2 bound in R5020 stimulated conditions (GRHL2 gain). B) Heatmaps showing CUT&RUN signal of GRHL2 in untreated and 4 hour 1 nM R5020 stimulated conditions, and 4 hour 1 nM R5020 stimulated PR at PR-only, GRHL2 bound in R5020 stimulated conditions and GRHL2 R5020 stimulated and PR co-occupied CUT&RUN peaks. C) Top 5 transcription factor motifs that were determined as enriched in all untreated GRHL2 peaks, all R5020 treated GRHL2 peaks, all R5020 treated PR CUT&RUN peaks, and all GRHL2 R5020 treated and PR co-occupied CUT&RUN peaks as determined by HOMER motif analysis^40^.

When we plotted the CUT&RUN signal on the ChIP-seq called peaks identified in Fig2 (**SupFig1C**), we found that the CUT&RUN GRHL2-signal increased upon R5020 treatment. In contrast, the CUT&RUN GRHL2 signal observed under untreated conditions, shows substantial overlap with GRHL2 and GRHL2–PR co-bound ChIP-seq peaks. This suggests that the R5020-dependent GRHL2 binding pattern observed in our CUT&RUN analysis mirrors the reported GRHL2 ChIP-seq profiles in Fig2 (**SupFig1C**).

Taken together, these results show that PR and GRHL2 can bind to the DNA independently, but also frequently overlap at enhancer regions. Moreover, GRHL2 chromatin binding is redistributed upon progesterone stimulation. This supports a model in which PR and GRHL2 cooperate to orchestrate hormone-dependent transcriptional regulation.

### GRHL2 and PR co-regulate a subset of genes involved in breast development

Next, we sought to identify transcriptional programs regulated by GRHL2 and PR. To this end, we generated T47DS cells with a stable shRNA-mediated knockdown of *GRHL2* (**SupFig2A-B**) and performed RNA-sequencing on wild-type and *GRHL2* knockdown T47DS cells treated with either vehicle or R5020 for 24 hours. Principal component analysis (PCA) showed that 97% of the variance in gene expression was explained by the R5020 stimulation (PC1) or *GRHL2* knockdown (PC2), confirming robust transcriptional responses to our experimental conditions (**SupFig2C**). To identify GRHL2 and PR co-regulated genes, we generated a stringent selection based on three criteria: 1) statistically significant differential gene expression after R5020 treatment, 2) statistically significant differential gene expression in GRHL2 knockdown cells, and 3) a significantly altered transcriptional response to R5020 upon GRHL2 depletion (**Fig4A**). Applying these criteria, we identified 549 genes regulated by both PR and GRHL2. The majority (70%) of these genes was repressed in response to R5020 stimulation (**Fig4B**). Gene Ontology enrichment analysis identified genes involved in breast tissue development, as well as cytoskeletal and cilium organization and assembly (**Fig4C**). We confirmed that 144 of these 549 co-regulated genes (∼26%) also exhibited at least a two-fold expression change as early as 4 hours post R5020 treatment (**SupFig2D**), suggesting that at least a subset of these co-regulated genes are direct PR targets. We further validated the expression changes by qRT-PCR for selected target genes, including *GRHL2* itself, *IGFBP5* and *TGFB2* (**Fig4D**, **SupFig2C-D**). Together, these results reveal a large number of genes that are jointly regulated by GRHL2 and PR, confirming their co-regulatory role in hormone responsive breast cancer cells.

**Fig 4.**
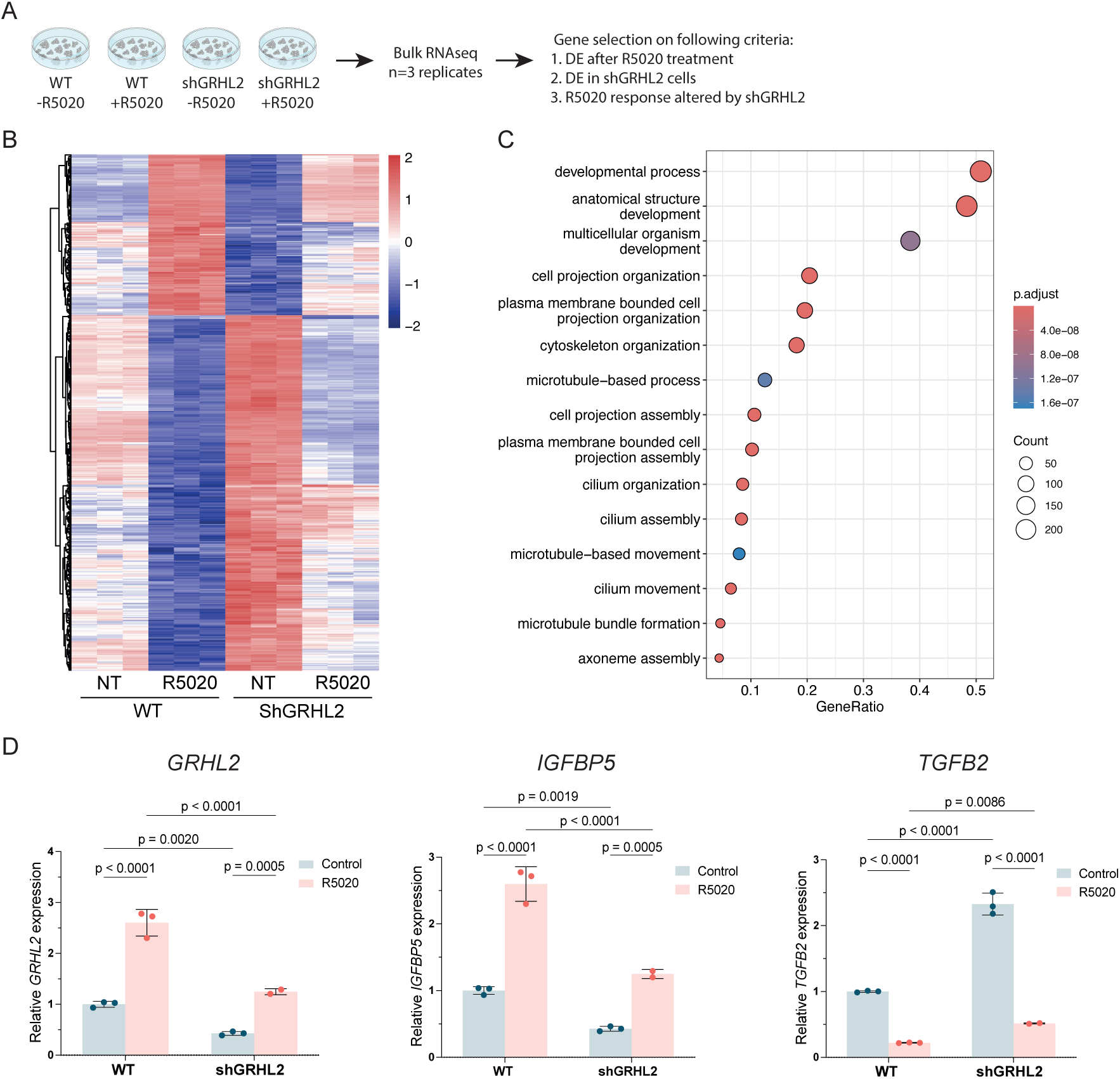
GRHL2 and PR co-regulate target genes involved in breast development and cilium organization. A) Schematic overview of the bulk RNA-seq process on wildtype (WT) T47DS cells and shRNA mediated *GRHL2* knockdown (shGRHL2) T47DS cells stimulated with 1 nM R5020 for 24 hours and the criteria used to select GRHL2 and PR co-regulated genes. B) Heatmap of bulk RNA-seq data from WT and shGRHL2 T47DS cells stimulated with 1 nM R5020 for 24 hours. Heatmap shows unsupervised clustering and expression changes of 549 genes that were selected based on the criteria listed in (A). Expression values are depicted as z-scores for n=3 replicates. NT= non-treated stripped-serum condition. C) Dot plot showing Go term enrichment analysis on the 549 selected GRHL2 and PR co-regulated genes. Enrichment analysis and figure generation was done using Clusterprofiler^41^. D) Bar graphs of qRT-PCR data, depicting the mean relative expression of *GRHL2, IGFBP5* and *TGFB2* after 4 hours of 1 nM R5020 treatment in WT or shGRHL2 T47DS cells. Reference gene: *YWHAZ*. Datapoints: individual values for n=2/3 biological replicates depicted as mean fold change normalized to control. P-values were calculated using a two-way ANOVA followed by an Uncorrected Fisher’s Least Significant Difference test.

### Chromatin looping connects GRHL2 and PR sites to distal target genes

Despite significant advances in our understanding of 3D genome organization, functional linking of enhancers to their target genes remains challenging^42^. In most studies, enhancers are assigned to the nearest gene, despite that fact that that 33– 73% of enhancers do not necessarily regulate their closest gene^43–45^. This is important since PR, as well as many other transcription factors, frequently bind distal enhancers located between 30 kb and 1 Mb away from their true associated TSSs^46,47^. High-resolution chromatin conformation capture techniques—such as Hi-C, 5C, ChIA-PET, and HiChIP—enable the identification of long-range chromatin interactions that are essential for transcriptional regulation by such factors^42^.

With this in mind, we sought to physically link the genomic GRHL2 and PR sites (as identified by ChIP-seq, **Fig2**), to the genes they regulate (as identified by RNA-seq, **Fig4**). We reasoned that if a GRHL2 or PR site was involved in the regulation of a specific gene, it should loop to the TSSs of this gene. To test this hypothesis, we used a previously published PR HiChIP dataset obtained using T47D cells^36^. HiChIP is an extension of Hi-C that captures protein-specific chromatin loops by enriching for DNA interactions bound by a transcription factor or histone modification of interest. Each chromatin loop contains two anchors, one on either end, representing interacting DNA regions brought into proximity by 3D chromatin looping. The PR HiChIP captured 7,076 chromatin loops involving PR without stimulation and 9,591 chromatin loops involving PR at one or both anchors after 30 minutes of R5020 stimulation (**Fig5A)** Only 987 of those loops overlapped, showing substantial rewiring of the PR mediated chromatin loops upon R5020 stimulation, as early as 30 minutes after stimulation (**Fig5A)**.

**Fig 5.**
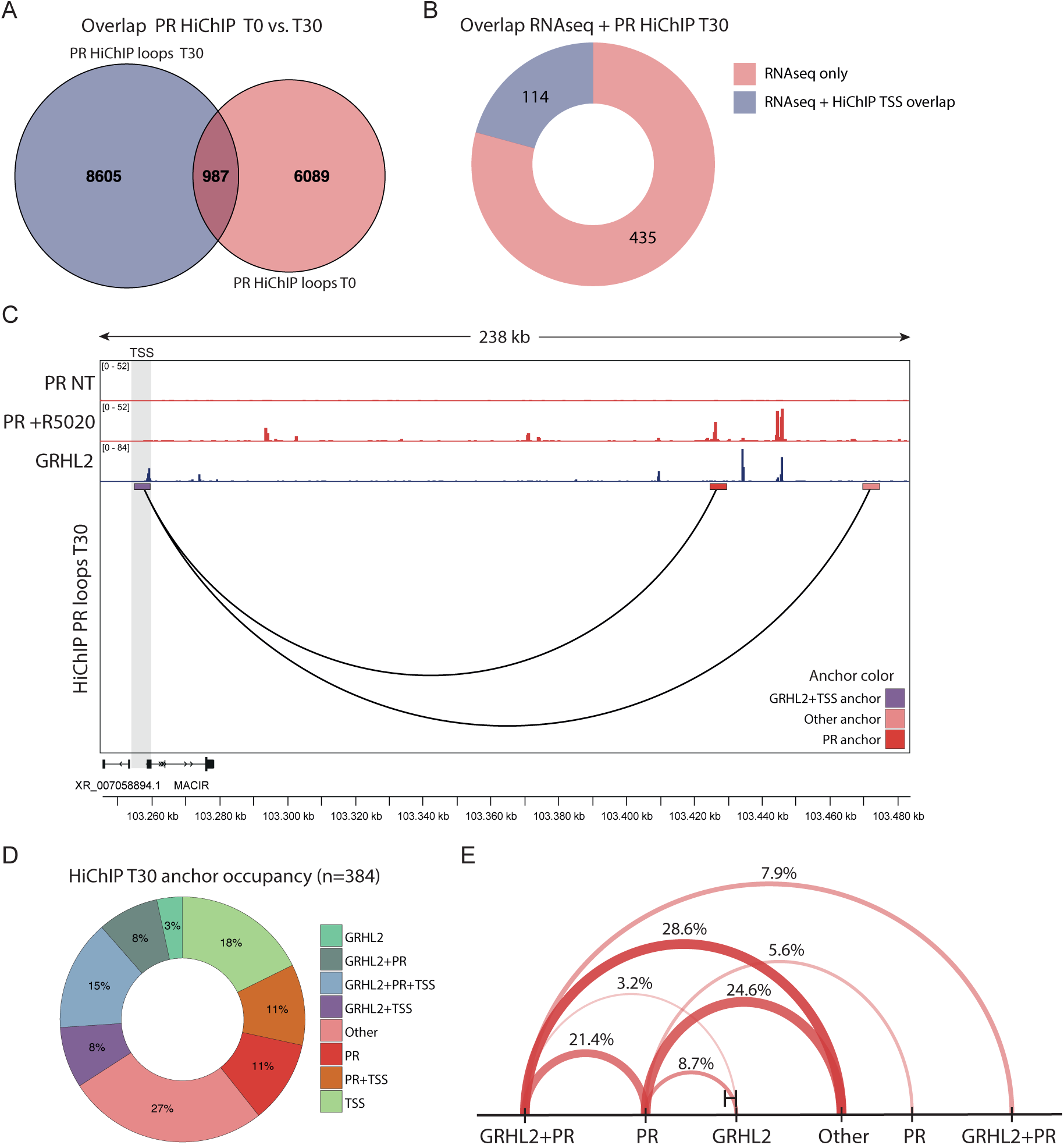
Integrative analysis of HiChIP, ChIP-seq, and RNA-seq reveals functional GRHL2–PR loops. A) Venn diagram showing the overlap between PR HiChIP loops in the non-treated (T0) condition and the 30 minutes 10 nM R5020 treated condition (T30). Replotting of data (i.e. number of called loops from Zaurin *et* al.^36^ B) Donut chart exhibiting the proportion of transcription start sites (TSSs) of the GRHL2 and PR regulated genes that overlaps with an anchor from a previously published PR HiChIP^36^. Of the 549 GRHL2 and PR regulated genes identified by RNA-seq, 114 genes have a TSS that overlaps with a HiChIP anchor. The TSS of the remaining 435 genes does not overlap with a HiChIP anchor. C) Visual representation of how GRHL2 and PR peaks converge on the *MACIR* gene promotor by chromatin looping, showing PR ChIP-seq^36^ peaks, GRHL2 ChIP-seq^23^ peaks and PR HiChIP loops^36^ directly anchoring to the GRHL2/PR regulated TSS. Anchors were colored by GRHL2 and PR occupancy as depicted in the legend at the lower right corner, anchors without GRHL2 or PR ChIP-seq signal or an annotated TSS were defined as ‘Other’. The gene TSS is highlighted with the grey bar. Data was visualized in the IGV browser^37^. D) Donut chart illustrating the HiChIP^36^ anchor occupancy categorized as overlapping with GRHL2 peak, PR peak, TSS, combinations of these three or other. The 384 included anchors (two for each of the 192 loops) were associated with the TSS of the 114 GRHL2 and PR regulated genes that overlap with a HiChIP^36^ anchor. E) Visual representation of the frequency of anchors defined as overlapping with GRHL2, PR, GRHL2+PR ChIP-seq peaks or other at opposing anchor pairs of the 192 selected HiChIP^36^ loops. The 126 loops were selected to have a PR ChIP-seq peak on at least one of the anchors. Color and thickness of the line represent the occurrence frequency.

Next, we integrated the PR HiChIP data with the GRHL2 and PR ChIP-seq and RNA-seq data. Here, we utilized the ChIP-seq data instead of our CUT&RUN dataset, as HiChIP is more methodologically similar to ChIP, and therefore more likely to integrate effectively. First, we took all 549 GRHL2 – PR co-regulated genes from our RNA-seq analysis and assessed if their TSSs overlapped with an anchor of at least one of the HiChIP loops. Using this approach, we identified 114 genes that are transcriptionally regulated by GRHL2 and PR and that also harbor a PR HiChIP loop anchor at or near their TSS (**Fig5B**). In total, we identified 192 loops directly connecting to the 114 GRHL2 and PR regulated genes (**Fig5B-C**). These loops extended up to 750kb, with the majority spanning 50-100kb (**SupFig3A**). We subsequently wondered if and to what extent GRHL2 and PR could be detected at the anchors of these 192 loops. Therefore, we aligned the GRHL2 and PR ChIP-seq data to assess their occupancy at each anchor. For all 192 loops, we categorized the associated 384 anchors according to the presence of a GRHL2 and/or PR ChIP-seq peak and/or a TSS (**Fig5C**,**D**). Among the annotated anchors, 23% were shared between GRHL2 and PR (∼two-fold enrichment compared to the entire ChIP-seq dataset, **Fig2B**), of which 15% was located at or near a TSS. Another 22% only contained a PR peak, of which 11% located at or near a TSS, while 11% only contained a GRHL2 peak, of which 3% located at or near a TSS (**Fig5D**). Notably, 27% of loop anchors, fell into the ‘Other’ category, meaning that they lacked either GRHL2, PR, or a TSS annotation (**Fig5D**). We hypothesize that these anchors are likely occupied by additional transcriptional regulators involved in the joint control of GRHL2 and PR target genes. To explore this possibility, we performed motif enrichment analysis specifically at these sites (**SupFig3B**). This analysis revealed no transcription factors that have specifically been linked to the breast before. However, we observed significant enrichment CTCFL and CTCF (outside the top 5), indicating that CTCF-mediated chromatin looping potentially contributes to the 3D regulatory context in which GRHL2 and PR exert their function (**SupFig3B**).

To further investigate the cooperative function of GRHL2 and PR in gene regulation, we analyzed their occupancy at opposing anchors of the PR HiChIP loops (**Fig5E**). Here, we only included loops that have a PR ChIP-seq peak on at least one of the two anchors, resulting in a selection of 126 loops. Of these, ∼61% contain at least one anchor that contains both GRHL2 and PR ChIP-seq peaks, indicating that the most prevalent mode of GRHL2–PR–mediated gene regulation potentially involves co-binding of GRHL2 and PR to at least one regulatory element (**Fig5E**). Nonetheless, all other combinations of GRHL2 and PR occupancy were also observed, suggesting that distinct GRHL2- and PR-bound elements can converge through 3D chromatin architecture to jointly regulate gene expression (**Fig5E**, **SupFig3C**). Moreover, some PR, GRHL2 or GRHL2-PR overlapping peaks not only establish direct connections with gene promoters but also interact with additional GRHL2 and PR (as well as other transcription factor) binding sites (**SupFig3C**).

### An integrative model for GRHL2 and PR transcriptional coregulation

CTCF- and Cohesin-mediated looping is well established as a key mechanism facilitating long-range enhancer–promoter interactions^48^. In our RIME, we identified GRHL2 to be associated with multiple Cohesin complex components and related regulatory factors (**SupFig4A**). Furthermore, publicly available ChIP-seq datasets for CTCF^49^ and RAD21^49^ in non-treated and 30 minutes R5020 treated conditions revealed that while CTCF binding is relatively stable, RAD21 binding to chromatin is highly enriched upon R5020 treatment (**SupFig4B-C**). Since TAD boundaries are relatively stable irrespective of R5020 presence^50^, we hypothesize that a subset of GRHL2/PR/TSS interactions are mediated by CTCF/Cohesin mediated looping, thus playing a role in the induced spatial proximity between GRHL2–PR enhancers and their target promoters. Furthermore, we note that RAD21/ the Cohesin complex is possibly recruited to the chromatin together with GRHL2 upon progesterone presence.

Taking all data together, we propose a model in which, in the absence of hormonal stimulation, GRHL2 and PR are already present in the nucleus and capable of physically interacting with each other (**Fig6**, **left**). Under these conditions, PR is largely unbound to its genomic target sites, whereas GRHL2 is bound to regions not associated with GRHL2 – PR target genes, as well as dynamically and at much lower levels, to shared GRHL2/PR sites (**Fig6**, **left**). Most genomic target sites and regulatory elements bound by either or both factors are not yet in the proximity of the promoter of their respective target gene (**Fig6**, **left**). In the presence of progesterone or, in an experimental setting, upon R5020 stimulation, the levels of nuclear GRHL2 and PR subtly increase, and transcriptionally active PR is recruited to GRHL2/PR shared and PR only binding sites. At the same time, GRHL2 chromatin binding is strengthened and reorganized upon progesterone stimulation. Together, PR and GRHL2 thus trigger extensive chromatin reorganization.

**Fig 6.**
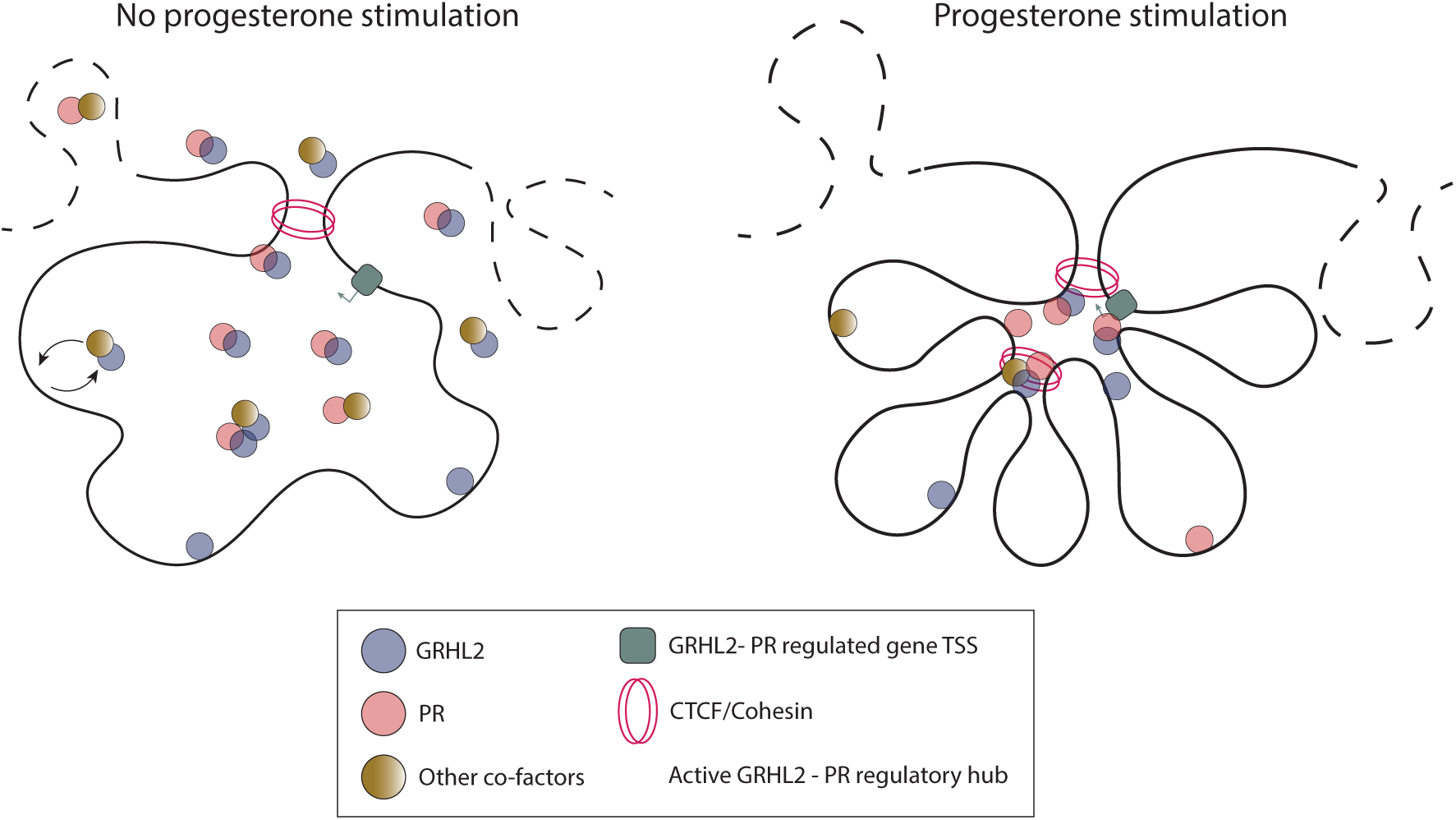
A model for gene expression regulation by GRHL2 and PR. Model for the co-regulatory function of GRHL2 and PR in target gene regulation. In absence of hormone (left), GRHL2 and PR form a complex, GRHL2 is dynamically bound to regions not associated with GRHL2 – PR target genes, as well as dynamically and at much lower levels, to shared GRHL2/PR sites, PR is largely not chromatin bound. As a result, GRHL2 and PR do not interact with their target promotors. Progesterone stimulation (right) triggers the recruitment of GRHL2 and PR to enhancers and the formation of both CTCF/Cohesin-dependent and independent chromatin loops to connect distal elements co- or individually bound by GRHL2, PR or co-factors to their target genes.

In addition to binding each other, GRHL2 and PR also associate with various co-factors. This results in locally high concentrations of GRHL2 and PR, facilitated by both CTCF/Cohesin-dependent and -independent loops to connect distal enhancers to target promoters (**Fig6**, **right**). While more than half of all PR-looped enhancers are co-bound by both GRHL2 and PR on at least one anchor, distinct GRHL2- or PR-bound elements also converge on promoters (**Fig6 right**). Moreover, multiple enhancer sites—bound by GRHL2 and/or PR as well as other transcriptional co-factors—are brought in close proximity with gene promoters through direct and potentially by indirect looping, enabling efficient transcription of genes involved in progesterone-driven breast tissue development and physiology (**Fig6**, **right**).

## Discussion

In this study, we provide a detailed mechanistic framework for the coordinated transcriptional regulation by the pioneering transcription factor GRHL2 and the steroid hormone receptor PR. By integrating proteomic, transcriptomic, and genomic analyses in T47D and T47DS cells in both unstimulated and R5020 stimulated conditions, we propose a model in which, in the absence of hormone, GRHL2 and PR form a pre-assembled nuclear complex that is associated with other chromatin modifying factors (**Fig1A-D** and **Fig6**, **left**). GRHL2 and PR dependent binding sites are mainly unlooped and spatially distant from their promoters. Upon progesterone stimulation, GRHL2 and PR are recruited to and/or enriched at specific enhancers, initiating hormone-dependent chromatin looping, that brings distal regulatory elements into contact with their target gene promoters (**Fig6**, **right**). These elements may be co-bound by GRHL2 and PR or individually bound by either factor or co-factors, together activating transcriptional programs essential for breast development and physiology. Thus, while the overall nuclear GRHL2 interactome does not undergo major changes upon hormone stimulation (**Fig1E**), these findings uncover a dynamic, hormone-responsive regulatory architecture that enables precise spatiotemporal control of gene expression in breast cells.

Using RIME and co-immunoprecipitation, we identified PR as a progesterone-independent interactor of GRHL2 under serum-stripped conditions. A study by Mohammed *et al*.^35^ also identified the GRHL2–PR interaction using PR RIME in both MCF7 and T47D cells, but reported it as progesterone-dependent. Although both studies detect an interaction between PR and GRHL2, our findings differ in the hormonal dependency of this association. Specifically, Mohammed *et al*.^35^ employed a non-treated condition involving full-serum medium, which contains endogenous estrogen and progesterone and can mimic physiological hormone stimulation. In contrast, our non-progesterone-stimulated condition was conducted under stripped-serum conditions, thus depleted of both estrogen and progesterone. Since, GRHL2 has been shown to associate with ER and to play a role in estrogen receptor dynamics and target gene expression^23,25,51^, suggesting that GRHL2 functions in both estrogen and progesterone signaling pathways. It is therefore plausible that, in the absence of both estrogen and progesterone, GRHL2 preferentially binds PR, and that this interaction is maintained, or potentially enhanced, upon progesterone stimulation. Conversely, in the presence of estrogen, either alone or together with low levels of progesterone, GRHL2 may associate more strongly with ER than with PR. Together, this raises important questions regarding how GRHL2’s DNA binding activity and interactome are regulated under fluctuating hormonal conditions, and how such modulation contributes to transcriptional programs in hormone-responsive cells.

Among the ten GRHL2 interactors identified as progesterone dependent in our RIME, most lacked direct links to GRHL2 or hormone signaling in the literature. However, STAT6 is co-expressed with PR in breast cancer and plays a role in luminal mammary epithelial cell development during pregnancy^52,53^. Our data indicates that the GRHL2– STAT6 interaction is lost upon progesterone stimulation, suggesting that STAT6 is either released from the DNA or redirected to other progesterone-responsive genes. The latter is supported by the fact that we found STAT6 motifs to be enriched in the ’Other’ (i.e. non-GRHL2 bound, non-PR bound, non-TSS) loop anchors connected to the GRHL2 and PR regulated genes. Therefore, it will be interesting to explore the exact role of STAT6 in GRHL2 and PR mediated gene regulation. In addition, the lysine methyltransferase SMYD2, known to methylate both H3K4 and non-histone proteins^54^, emerged as a progesterone dependent GRHL2 interactor. Notably, SMYD2 directly methylates ER, modulating its transcriptional activity^55^. It is therefore plausible that, upon progesterone stimulation, SMYD2 is recruited to the GRHL2–PR complex, where it may methylate both PR and associated enhancer regions to fine-tune GRHL2–PR–mediated transcription, in a manner analogous to ER regulation.

In this study, we only briefly explored which additional transcription factors are involved in GRHL2 - PR mediated gene regulation. FOXA1 emerged as one of the top enriched motifs in a genome-wide analysis of GRHL2-bound, PR-bound, and GRHL2–PR co-bound sites (**Fig2F**), consistent with its known central role in mature luminal cells^56–59^. In support of this notion, we also detect FOXA1 as a GRHL2 binding protein in our RIME analysis. However, FOXA1 motif enrichment was not observed at the 994 loop anchors linked to the 114 genes jointly regulated by GRHL2 and PR, nor in the GRHL2 or PR peaks of our CUT&RUN. Interestingly, FOXA1 may compete with PR for enhancer occupancy, as it has been shown that FOXA1 depletion led to increased PR binding at enhancer sites^60^. These findings suggest that although GRHL2 and FOXA1 cooperate in some contexts, GRHL2–PR co-regulatory activity may occur largely independent of FOXA1.

Our CUT&RUN analysis showed that GRHL2 chromatin binding is redistributed and increased upon progesterone presence. This has not been reported before, but it is in line with GRHL2 being a direct PR target^61^ and with prior work showing that estrogen treatment leads to a genome-wide increase in GRHL2 binding^51^. Interestingly, we find that GRHL2 CUT&RUN peaks detected following R5020 treatment, but not those from untreated conditions, overlap with GRHL2 sites identified by ChIP-seq under none-hormone stimulated conditions. This discrepancy may be explained by residual hormone activity during the original ChIP-seq experiment, or by methodological differences: unlike ChIP-seq, CUT&RUN does not involve cross-linking and is therefore more likely to capture stable DNA–protein interactions. Transient or weaker interactions, such as those involving dynamic transcription factors like GRHL2 and PR, may only be detectable by CUT&RUN when stabilized (e.g. by hormonal treatment), whereas ChIP-seq may capture these interactions more readily. These methodological and biological differences may also explain why the number of GRHL2, and PR peaks identified in our CUT&RUN data is nearly two orders of magnitude lower than in the reanalyzed ChIP-seq dataset (see **Fig3B** vs. **Fig2B**).

Ultimately, it remains challenging to determine the precise function of individual enhancers and to accurately link them to their target genes. This difficulty largely stems from the location of enhancers, which are often at considerable distance from the genes they regulate, as well as from their ability to bypass nearby, and interspersed genes. Nonetheless, mapping enhancer–target gene relationships is crucial for understanding enhancer function in both normal biology and disease. In the absence of individual experimental validation, chromatin conformation-based approaches currently offer the most effective way to predict enhancer–promoter interactions. Therefore, we utilized PR HiChIP data to link GRHL2 and/or PR sites to their target genes. While this method provides higher confidence in target assignment than assigning peaks to the nearest gene, it is not without limitations. Notably, the PR HiChIP dataset we used was generated at an early timepoint following progesterone stimulation (i.e. 30 minutes). Although this enabled the identification of the earliest GRHL2–PR co-regulated targets, additional timepoints would capture a broader dynamic range of interactions, increase the number of assignable targets, and provide insights into the temporal dynamics of PR-mediated enhancer–promoter looping. Moreover, since this dataset is PR-centered, it does not capture GRHL2-mediated chromatin loops that do not directly involve PR but may converge on the same promoters. As a result, we may be underestimating the extent of GRHL2’s regulatory influence and the full landscape of GRHL2–PR co-regulated target genes. However, separate PR and GRHL2 HiChIP datasets are also not ideal as they do not allow direct detection of PR and GRHL2 overlapping anchors. Unfortunately, a method to measure the protein directed genome architecture of two transcription factors has not been developed yet.

In summary, this study expands our knowledge on the regulatory landscape of progesterone signaling in the breast epithelium by integrating genomic, transcriptomic and proteomic data. We were able to uncover a previously unrecognized partnership between GRHL2 and PR that orchestrates hormone-responsive gene expression through dynamic chromatin binding and 3D genome organization. Our model poses that GRHL2 and PR form a complex in the absence of hormone but under these conditions, do not stably bind chromatin. Progesterone stimulation recruits GRHL2-PR complexes to enhancers followed by the formation of chromatin loops, that connect distal elements to target gene promoters. Co-binding or individual binding by GRHL2, PR, or co-factors then activates key transcriptional programs. This model provides a broad framework for understanding GRHL2–PR-mediated gene regulation. However, we find that each individual target gene ultimately is regulated by a unique combination of distal elements individually or jointly bound by GRHL2, PR and other transcription factors, which loop to its promoter. Our integrated analysis highlights the intricate and gene-specific nature of these regulatory interactions, illustrating the complexity that can arise even from the interplay of just two transcription factors.

## Materials & methods

### Cell culture

Human T47DS breast cancer cells (a kind gift from Dr. Stieneke van den Brink, Hubrecht institute, Utrecht, The Netherlands) and HEK293TN cells (System Biosciences, #LV900A-1) were routinely cultured in Dulbecco’s Modified Eagle Medium (DMEM) containing GlutaMAX (Gibco, #11584516), supplemented with 10% Fetal Bovine Serum (FBS) (Thermo Fisher Scientific, #11573397). Cells were split 1:10 every 3-4 days and routinely tested for mycoplasma. When treated, T47DS cells were cultured in DMEM containing GlutaMAX (Gibco, #11584516), supplemented with 5% Charcoal Stripped Fetal Bovine Serum (Thermo Fisher Scientific, #A3382101) for 24 hours, followed by stimulation with 1 nM R5020 (Perkin Elmer, #NLP004005MG) for the indicated time, again in stripped-serum conditions.

### Generation of T47DS GRHL2 knockdown lines

For lentiviral production, 5 × 10⁶ HEK293TN cells were seeded in 10 cm dishes. The following day, cells were transfected with 3 µg packaging vector pCMVDR8.2 (Addgene, #8455), 3 µg RSV-rev (Addgene, #12253), 3 µg VSV-G (Addgene, #12259), and 8 µg pLKO.1 GRHL2 shRNA vector (RNAi Consortium, TRCN0000015812; a kind gift from Peter Stroeken, Amsterdam UMC). After 24 hours, the medium was refreshed. Viral supernatant was collected 48 hours post-transfection, filtered through a 0.45 µm filter, and diluted 1:2. T47DS cells were infected with the viral supernatant in the presence of polybrene (1:2,000; Merck Millipore, #TR-1003-G). Following 24 hours of incubation, cells were split and selected with puromycin (1 µg/mL, Thermo Fisher Scientific, #A1113803) for up to two weeks to generate stable GRHL2 knockdown lines.

### Rapid IP-mass spectrometry of endogenous protein (RIME)

RIME analysis was performed as previously described by Mohammed *et al.*^31^. Briefly, 6x10^7^ T47DS per condition were crosslinked for 8 minutes by adding formaldehyde (Thermo Fisher Scientific, #28908) to the medium to a final concentration of 1%. The reaction was quenched by adding Glycine to a final concentration of 0.1 M. The cells were washed twice with ice-cold PBS supplemented with protease inhibitors (Sigma-Aldrich, #P8340, 1:500) and collected by scraping them off the plate. Cells were then pelleted by centrifugation 2,000 g for 3 minutes at 4 °C and lysed rotating for 5 min at 4 °C using lysis buffer LB1 (50 mM HEPES-KOH, (pH 7.5), 140 mM NaCl, 1 mM EDTA, 10% (vol/vol) glycerol, 0.5% (vol/vol) NP-40 and 0.25% (vol/vol) Triton X-100). This was followed by a second lysis in LB2 (10 mM Tris-HCL (pH 8.0), 200 mM NaCl, 1 mM EDTA and 0.5 mM EGTA), again while rotating for 5 min at 4 °C. The sample was then resuspended in LB3 (10 mM Tris-HCl (pH 8.0), 100 mM NaCl, 1 mM EDTA, 0.5 mM EGTA, 0.1% (wt/vol) sodium deoxycholate and 0.5% (vol/vol) *N*-lauroylsarcosine) and divided into three separate Eppendorf tubes for sonication in a Bioruptor pico. (Diagenode) at 4 °C using a 30 seconds on/off cycle for 10 cycles. The sonicated samples were pooled again, Triton X-100 was added to a final volume of 1% and the lysate was cleared by centrifugation. In the meantime, Dynabeads (Protein A, Invitrogen, #10001D) were washed four times in PBS containing 5 mg/ml BSA. 5 mg GRHL2 antibody (Sigma Aldrich, #HPA004820) and IgG (Santa cruz, #sc-2027X, as a negative control) were conjugated to the beads in PBS/BSA solution for 1 hour at room temperature. Sonicated nuclear lysates were pooled with antibody conjugated beads and incubated rotating at 4 °C overnight. The next day, beads were washed with RIPA buffer (50 mM HEPES (pH 7.6), 1 mM EDTA, 0.7% (wt/vol) sodium deoxycholate, 1% (vol/vol) NP-40 and 0.5 M LiCl) for a total of ten washes. Three technical replicates were performed for each experimental condition.

From this point forward processing and analysis was performed by the Laboratory for Mass spectrometry of Biomolecules, Swammerdam Institute for life sciences, University of Amsterdam. In brief, input samples were processed as described in Hughes *et al.*^62^. Protein concentration was quantified using the Pierce BCA protein assay (Thermo Fisher Scientific, #23227), following the manufacturer’s protocol and all samples were reduced and alkylated in one step. Pull-down samples were digested on-bead according to Mohammed *et al.*^31^. Peptide samples were analyzed using an Ultimate 3000 RSLCnano UHPLC system (Thermo Fisher Scientific, Germeringen, Germany) coupled to a TIMS-TOF Pro mass spectrometer (Bruker, Bremen, Germany). Raw data were processed using MaxQuant (version 2.6.2.0), with trypsin/P specified as the digestion enzyme allowing up to two missed cleavages. Carbamidomethylation of cysteine was set as a fixed modification, and oxidation of methionine as a variable modification. Spectra were searched against the human Uniprot proteome database. MaxQuant IBAQ output values were used for downstream analysis using Perseus (version 2.1.3.0). Potential contaminants, proteins that were only identified by peptides that carry one or more modified amino acids, and proteins identified from a decoy sequence were filtered out. Values were Log2 transformed and rows with more than two out of three missing values per experimental group were discarded. Other NaN values were replaced with 1 and the three technical replicates were used to identify differentially expressed proteins between conditions using a two-sample Student’s t-test with a permutation-based false discovery rate (FDR) correction to account for multiple testing. A significance threshold of FDR < 0.05 was applied. Raw and processed RIME data are publicly available via: https://osf.io/9uqsw/.

### Co-immunoprecipitation

Co-immunoprecipitation of GRHL2 in T47DS cells was performed using approximately 3×10⁸ cells. Cells were washed once with PBS containing protease inhibitors (Sigma-Aldrich, #P8340, 1:500) directly on the dish, then harvested by scraping. Pelleted cells were resuspended in lysis buffer (50 mM Tris-HCl (pH 7.40), 5 mM EDTA, 0.25% Triton X-100, 10% glycerol, 100 mM NaCl) and incubated for 30 minutes at 4 °C with gentle rotation. Lysates were cleared by centrifugation at 2,000 × g for 20 minutes at 4 °C. An aliquot of the supernatant was reserved as input for the whole lysate control.

The remaining lysate was incubated overnight at 4 °C with 5 µg GRHL2 antibody (Sigma-Aldrich, #HPA004820) under rotation. The following morning, 50 µL of pre-equilibrated Dynabeads Protein A (Invitrogen, #10001D) was added, and samples were rotated for 1 hour at 4 °C. Beads were washed three times with lysis buffer, and bound proteins were eluted by boiling in elution buffer (50% lysis buffer, 50% Laemmli buffer; Sigma-Aldrich, #38733) at 95 °C for 5 minutes. Eluted proteins were subsequently analyzed by western blot.

### Western Blot

Western blotting was performed as previously described^63^. In short, half of the immunoprecipitated or input samples were loaded onto a 10% SDS-PAGE gel. Proteins were transferred to a 0.2 µm nitrocellulose membrane using the Trans-Blot Turbo Transfer System (Bio-Rad) and blocked with a 1:1 dilution of TBS and Odyssey Blocking Buffer (LI-COR Biosciences, #927-50100). Membranes were incubated overnight at 4 °C with primary antibodies diluted in blocking buffer supplemented with 0.1% Tween-20: anti-PR (1:1,000, Thermo Fisher Scientific, #MA5-16393, recognizing both PR-A and PR-B) and anti-GRHL2 (1:2,000, Sigma-Aldrich, #HPA004820). The following day, membranes were washed in TBS supplemented with 0.1% Tween-20 and incubated for 1 hour at RT with secondary antibodies diluted 1:20,000 in TBS with 0.1% Tween-20: IRDye 680LT (LI-COR, #926–68022) or IRDye 800CW (LI-COR, #926–32211). Signal detection was performed using the Odyssey Fc Imaging System (LI-COR Biosciences).

### RNA isolation, Library preparation and RNA-seq analysis

RNA isolation, processing, and analysis for RNA-seq were performed as described previously^61^. In summary, 700,000 T47DS WT or *GRHL2* knockdown cells (stable shRNA) were seeded in 6-well plates containing DMEM + GlutaMAX (Gibco, #11584516) supplemented with 5% charcoal-stripped fetal bovine serum (Thermo Fisher Scientific, #A3382101). 24 hours after plating, the medium was refreshed, and cells were treated for an additional 24 hours with either 1 nM R5020 or ethanol (vehicle control). RNA was isolated using the RNeasy Mini Kit (Qiagen, #74104) with on-column DNase digestion (Qiagen, RNase-Free DNase Set, #79254) according to the manufacturer’s instructions.

For library preparation, 1 µg of total RNA was subjected to poly(A) selection using the NEBNext Poly(A) mRNA Magnetic Isolation Module (New England Biolabs). RNA-seq libraries were generated using the NEBNext Ultra II Directional RNA Library Prep Kit and NEBNext Multiplex Oligos for Illumina (Unique Dual Index Primer Pairs; New England Biolabs), following the manufacturer’s protocols. Library size distribution was assessed using a 2200 TapeStation System with Agilent D1000 ScreenTapes (Agilent Technologies). Quantification was performed using the NEBNext Library Quant Kit for Illumina on a QuantStudio 3 Real-Time PCR System (Thermo Fisher Scientific). Libraries were sequenced (75 bp single end) using a NextSeq 500/550 High Output Kit v2.5 (75 Cycles) on a NextSeq 550 System (Illumina). All library preparation and sequencing was carried out by MAD: Dutch Genomics Service & Support Provider, Swammerdam Institute for Life Sciences, University of Amsterdam.

On the Galaxy server, raw FASTQ files were processed and quality control was performed using MultiQC^64^, followed by trimming of overrepresented sequences with Trimmomatic^65^ using a curated adapter list. Trimmed reads were aligned to the human reference genome (GRCh38.p14) using HISAT2^66^. Mapping quality was evaluated with SAMtools stats and summarized with MultiQC. Read counts were obtained using HTSeq-count^67^, and differential gene expression analysis was conducted in RStudio (version 2023.06.1) using DESeq2^68^. The data have been deposited with NCBI GEO under accession number GSE291778. A list of Log2fold changes of the 549 GRHL2 and PR co-regulated genes and the 114 genes that are transcriptionally regulated by GRHL2 and PR and harbor a PR HiChIP loop anchor at or near their TSS are available via: https://osf.io/9uqsw/. Go term enrichment on the selected PR and GRHL2 regulated genes was performed using the *enrichGO()* function of the Clusterprofiler R package^41^.

### RNA isolation and qRT-PCR

RNA isolation and qRT-PCR were performed as previously described^63^. Briefly, total RNA was extracted from T47DS cells using TRIzol Reagent (Invitrogen, #15596018) following the manufacturer’s protocol. Isolated RNA was treated with RQ1 DNase (Promega, #M6101) to remove genomic DNA contamination. cDNA synthesis was carried out using SuperScript IV Reverse Transcriptase (Invitrogen, #18090200) and Random Hexamers (Invitrogen, #N8080127), according to the manufacturer’s instructions, with the addition of RiboLock RNase Inhibitor (Thermo Fisher, #EO0328). Resulting cDNA samples were diluted 10-fold prior to qRT-PCR. Quantitative real-time PCR was performed using 5X HOT FIREPol EvaGreen qPCR Mix Plus (ROX) (Solis Biodyne, #08-24-00008) on a QuantStudio 3 Real-Time PCR System. Primer sequences are listed in Table 1. Relative expression levels were calculated using the ΔΔCt method and normalized to *YWHAZ* ^69^ and untreated control samples.

**Table 1.**
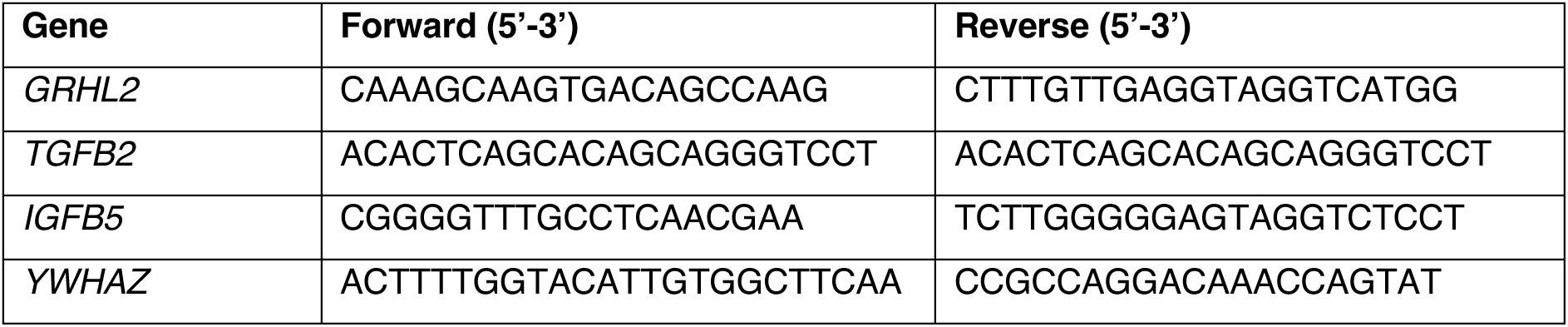
Primers used for qRT-PCR.

### CUT&RUN Low Volume Urea (LoV-U)

CUT&RUN LoV-U was performed as described in Zambanini *et al*.^70^, with the following modifications. Approximately 500,000 T47DS cells per sample were harvested using 0.25% trypsin, and cells were washed three times in nuclear extraction buffer (20 mM HEPES-KOH (pH 8.2), 10 mM KCl, 0.5 mM spermidine, 0.05% IGEPAL, 20% glycerol, and Roche Complete Protease Inhibitor EDTA-Free) to isolate nuclei. Nucleic pellets were flash-frozen in an isopropyl chamber and stored at –80 °C until further processing. Nuclei were thawed, washed once in nuclear extraction buffer and subsequently bound to equilibrated magnetic ConA agarose beads (Antibodies-online, #ABIN6952467). Following incubation, nuclei-bead complexes were washed for 5 minutes in wash buffer (20 mM HEPES (pH 7.5), 150 mM NaCl, 0.5 mM spermidine, and Roche Complete Protease Inhibitor EDTA-Free, 0.025% digitonin) supplemented with 2 mM EDTA, and then transferred to PCR tubes. Antibodies (anti-PR (1:100, Cell Signaling, #8757S), anti-GRHL2 (1:100, Sigma-Aldrich, #HPA004820), or rabbit IgG isotype control (1:100, Invitrogen, #100500C)) were added to the wash buffer and incubated overnight at 4 °C on a rotator. The next day, samples were washed five times with wash buffer, then resuspended in pAG-MN buffer (wash buffer containing 120 ng/sample pAG-MN) and incubated for 30 minutes at 4 °C on a rotator. Following another five washes, digestion was initiated by resuspending the beads in wash buffer supplemented with 2 mM CaCl₂ and incubated at room temperature for 5 minutes, followed by an additional 25 minutes at 4 °C. The reaction was stopped by adding 250 mM EDTA/EGTA mix. Fragment release was initiated by the addition of 5 M NaCl and incubation at 37 °C for 30 minutes. The supernatant was collected, and beads were subsequently resuspended in 1× Urea buffer (100 mM NaCl, 2 mM EDTA, 2 mM EGTA, 0.5% IGEPAL, 8.5 M urea) for a second release step. After 30 minutes at room temperature, the urea release was combined with the initial supernatant. Samples were treated with SDS and proteinase K, followed by phenol:chloroform:isoamyl alcohol extraction and ethanol precipitation. Libraries were sequenced on an Illumina NextSeq 550 platform using 36 bp paired-end reads to a depth of 5–15 million reads per sample.

### CUT&RUN Data Analysis

Quality of the reads was assessed using fastqc^71^. Reads were trimmed with bbmap bbdud^72^ to remove adapters and poly [AT], [G] and [C] repeat sequences. Reads were aligned to the hg38 genome with bowtie2^73^ with the options –local –very-sensitive-local –no-unal –no-mixed -no-discordant –phred33 –dovetail -I 0 -X 500. Samtools suite^74^ was used to remove duplicate and incorrectly paired reads. Bedtools^75^ was used to remove reads mapped to the CUT&RUN hg38 suspect list^76^ from bam files. Bedgraphs were created using Bedtools genomecov on pair-end mode. Peaks were called using MACS2^77^ with the options -f BAMPE and -p 1e-3 against the IgG control for narrowPeaks. The data will be deposited with NCBI GEO.

### Re-analysis of ChIP-seq datasets

Raw ChIP-seq datafiles from PR^36^ (Control; SRX11374768, and 1 nM R5020 treated; SRX11374766) and GRHL2^23^ (two replicates; SRX5350536, SRX5350537) datasets were downloaded from the SRA server. Quality of the reads was assessed using fastqc^71^ (version 0.11.9), followed by trimming of overrepresented sequences adapters and PCR primers with Trimmomatic^78^. Trimmed reads were aligned to the hg38 genome with bowtie2^73^ with the options –local –very-sensitive-local –no-unal –no-mixed -no-discordant – dovetail -I 0 -X 500. Samtools^74^ was used to convert sam to bam files, fix improperly paired mates, and remove duplicates. Bedtools^75^ was used to remove reads mapped to the hg38 blacklisted regions^79^. Final read counts of usable fragments were determined from filtered and deduplicated bam files. Bedgraphs were created with Bedtools genomecov on pair-end mode. Bedgraps were visualized in IGV^37^. For peak calling, visualization and signal graphs, replicate bam files were merged with SAMtools into a single file. Peaks were called using MACS2^77^ with the options -f BAMPE and -p 1e-3 for narrowPeaks. deepTools^80^ was used to convert bam files to normalized BigWig files (bamCoverage using -RPGC option and -e to extend reads to fragment length). Bigwigs were visualized in IGV; tracks shown in the figures are a merge of two biological replicates (GRHL2) or single replicate (PR), normalized to reads per genome coverage, and scaled by factor.

### Downstream analysis

For both ChIP-seq and CUT&RUN signal intensity heatmaps and signal profiles were generated using deepTools^80^. Peak overlaps were calculated using Bedtools^75^, requiring a minimum of 1 bp overlap. Venn diagrams were created with the *Venndiagram* R package, and genome annotation was performed using the *ChIPseeker* ^38,39^ R package. Motif enrichment analysis was carried out using HOMER^40^ with the options -size 200 and -mask. Processed PR HiChIP data from R5020-stimulated T47D cells (Zaurin *et al*.^36^, GEO accession: GSM5425945) were downloaded in .bedpe format and overlapped with TSS regions (–5 kb to +1 kb from the TSS) of 549 genes identified as PR- and GRHL2-regulated by RNA-seq, using Bedtools pairtobed^75^. TSS regions containing a HiChIP anchor were selected for further analysis. Peak-called ChIP-seq datasets for R5020-stimulated RAD21^49^ (SRX22674036) and CTCF^49^ (SRX22674025) were retrieved from ChIPatlas^81^.

## Statements and declarations

## Acknowledgements

We thank Gertjan Kramer and other members of the Laboratory for Mass spectrometry of Biomolecules, Swammerdam Institute for life sciences, University of Amsterdam for their support in Mass spectrometry analysis. We also thank Selina van Leeuwen and other members of the MAD: Dutch Genomics Service & Support Provider, Swammerdam Institute for Life Sciences, University of Amsterdam for their sequencing services. We thank our colleagues of the Developmental, Stem Cell & Cancer Biology (DSCCB) group and other colleagues in the Cell & Systems Biology theme at the Swammerdam Institute for Life Sciences for fruitful discussions and feedback during the project. We would like to thank all members of the Cantù lab for their guidance during our CUT&RUN experiments. We specifically thank our MSc internship student Jeltje Rommens for the initial analysis and workflow setup for identifying GRHL2 and PR regulated genes.

RvA acknowledges funding support from the Netherlands Organization for Scientific Research (OCENW.KLEIN.169 + NWO ALW VIDI 864.13.002). CC acknowledges support from Cancerfonden (21 1572 Pj and 24 3487 Pj 01 H), the Swedish Research Council, Vetenskapsrådet (2021–03075 and 2023-01898), Additional Ventures (SVRF2021-1048003), Linköping University, Joanna Cocozzas stiftelse för barnmedicinsk forskning. CC is a fellow of the Wallenberg Molecular Medicine (WCMM) and SciLifeLab and receive generous financial support from the Knut and Alice Wallenberg Foundation. MTA acknowledges funding support for a research visit to the lab of Claudio Cantù to perform CUT&RUN experiments from the Landelijk Netwerk voor Vrouwelijke Hoogleraren (LNVH) (Advancing Women in Biology fund 2024/2025) and the Genootschap ter bevordering van Natuur-, Genees- en Heelkunde.

## Author contributions

MTA and RvA conceived the study. AN and MTA designed, performed and analyzed the CUT&RUN experiments. MTA designed, performed and analyzed all other experiments. MTA performed bioinformatic data analysis using publicly available datasets. MTA conceived the GRHL2 - PR regulation model. RvA, ALvB and CC supervised the study. MTA wrote the manuscript with input from all authors.

## Declaration of the use of generative AI

No generative AI tools were used to generate content. During the writing process, ChatGPT was used to improve writing style of the first draft written by MTA. The output was critically reviewed and partly re-written by the author. The authors take full responsibility for the final text, content, claims and references in this work.

---

**Sup Fig 1.**
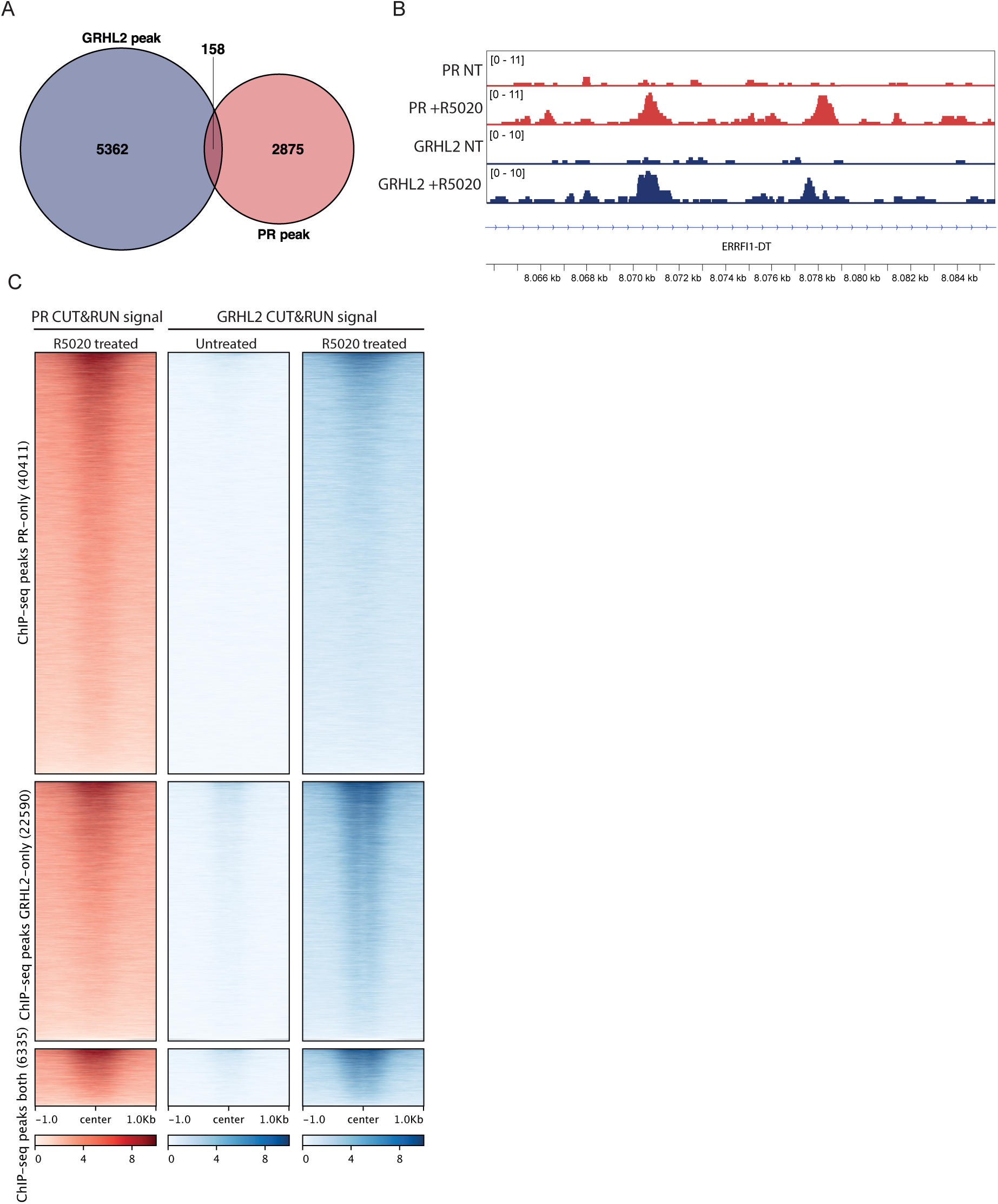
R5020-induced GRHL2 binding sites identified by CUT&RUN align with GRHL2 ChIP-seq profiles. A) Venn diagram showing the overlap between 4 hour 1 nm R5050 stimulated GRHL2 and PR CUT&RUN peaks. Peaks were considered overlapping if they overlapped by at least 1 bp. B) A representative example of two PR and GRHL2 overlapping CUT&RUN peaks. Data was visualized in the IGV browser^37^. NT= non-treated. C) Heatmaps showing CUT&RUN signal of PR R5020 treated and GRHL2 in untreated and 4 hour 1 nM R5020 stimulated conditions, at PR-only, GRHL2-only and GRHL2-PR shared ChIP-seq peaks.

**Sup Fig 2.**
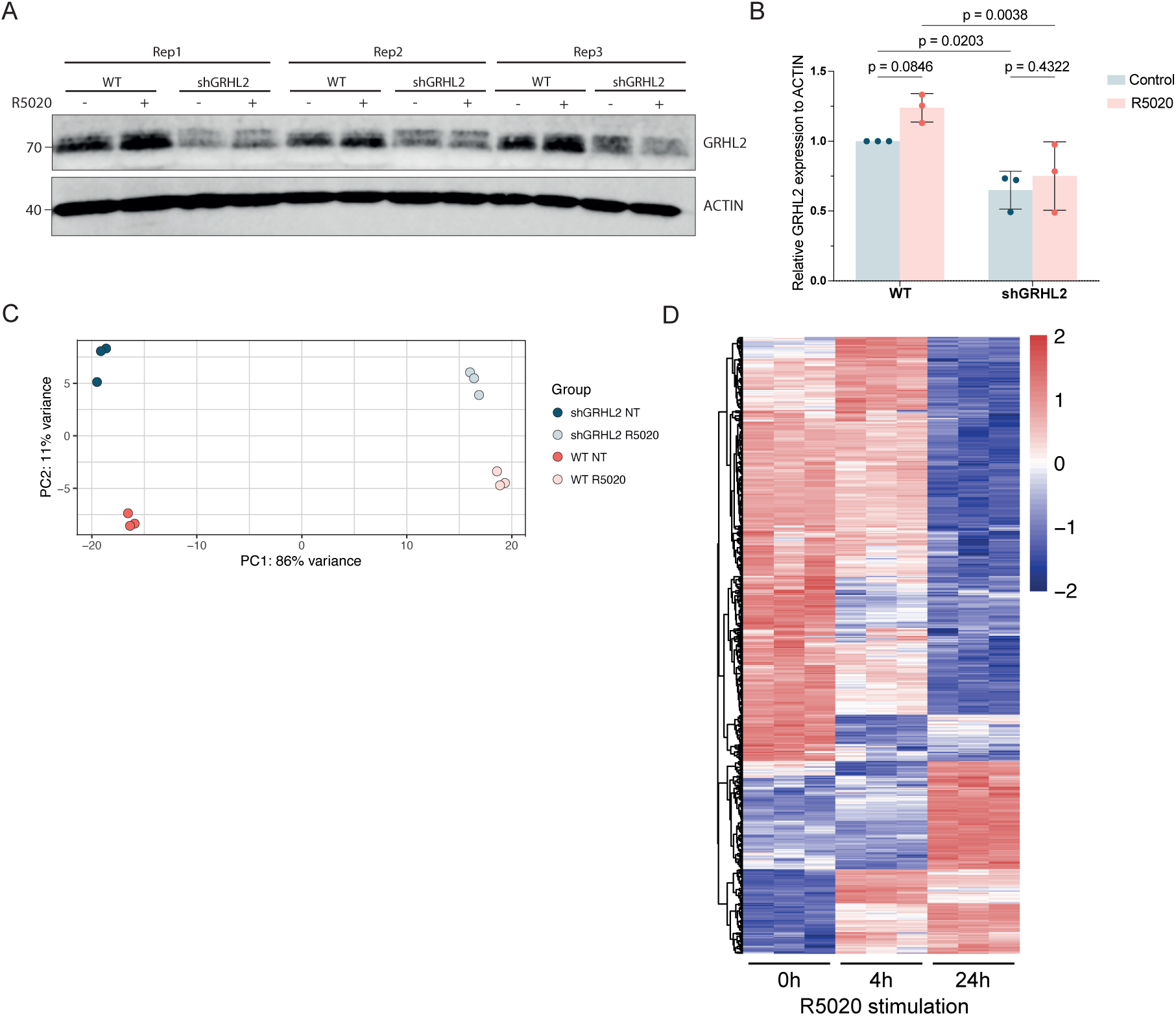
Validation of direct GRHL2 – PR target genes. A) Western blot for GRHL2 of wildtype (WT) and shRNA mediated *GRHL2* knockdown (shGRHL2) T47DS cells stimulated with 1 nM R5020 for 4 hours. Three independent biological replicates are shown. ACTIN was used as a loading control. B) Bar graph showing the relative GRHL2 protein expression over ACTIN of the western blot in (A). Individual values are normalized over the non-treated WT (control). Datapoints: individual values for n=3 biological replicates depicted as mean fold change normalized to control. P-values were calculated using a two-way ANOVA followed by a Uncorrected Fisher’s Least Significant Difference test. C) Principal component analysis (PCA) of all genes expressed in the bulk RNA-seq in WT and shGRHL2 T47DS cells that were non-treated (NT) or stimulated for 24 hours with 1 nM R5020. Each dot represents one replicate; each group contains n=3 replicates. D) Heatmap of bulk RNA-seq data from WT T47DS cells stimulated with 1 nM R5020 for 4 and 24 hours. Heatmap shows unsupervised clustering and expression changes of the 549 selected GRHL2 and PR co-regulated genes of n=3 biological replicates per condition.

**Sup Fig 3.**
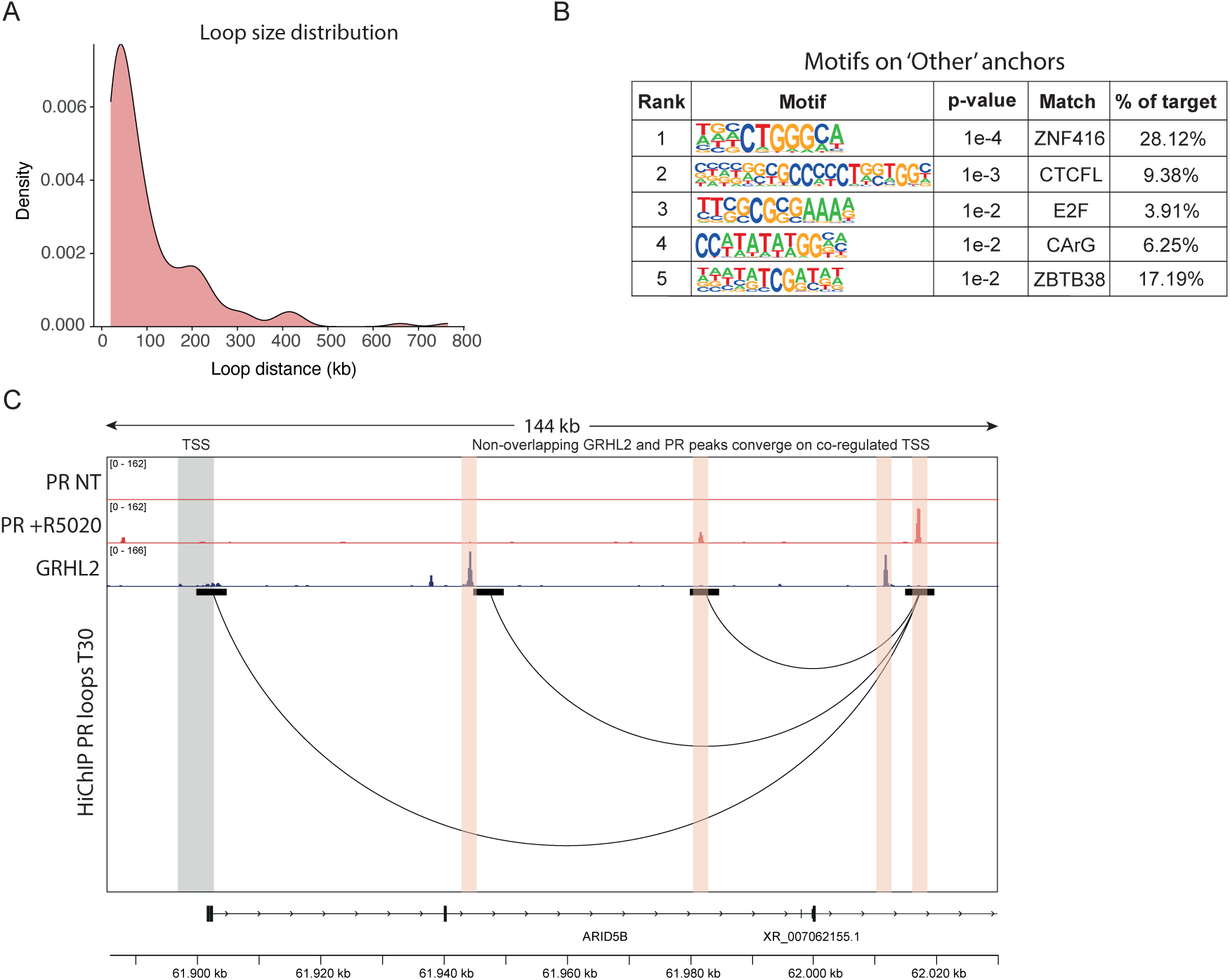
GRHL2- or PR bound enhancers can converge through 3D chromatin looping to jointly regulate gene expression. A) Density plot of the loop size distribution of the 192 PR HiChIP^36^ loops that are directly link to a GRHL2/PR regulated TSS as determined by RNA-seq. Loop size is displayed in kilobases (kb). B) Selection of transcription factor motifs that were determined as enriched at anchors that did not harbor a GRHL2 peak, PR peak, or regulated TSS, were defined as ‘Other’. Motif enrichment analysis was performed using HOMER^40^. C) Visual representation of individual non-overlapping GRHL2 and PR peaks, and the *ARID5B* promotor that converge on one PR peak by chromatin looping, showing PR ChIP-seq^36^ peaks, GRHL2 ChIP-seq^23^ peaks and PR HiChIP loops^36^. Grey bar highlights the *ARID5B* TSS. Orange bars highlight the individual PR and GRHL2 peaks. Data was visualized in the IGV browser^37^.

**Sup Fig 4.**
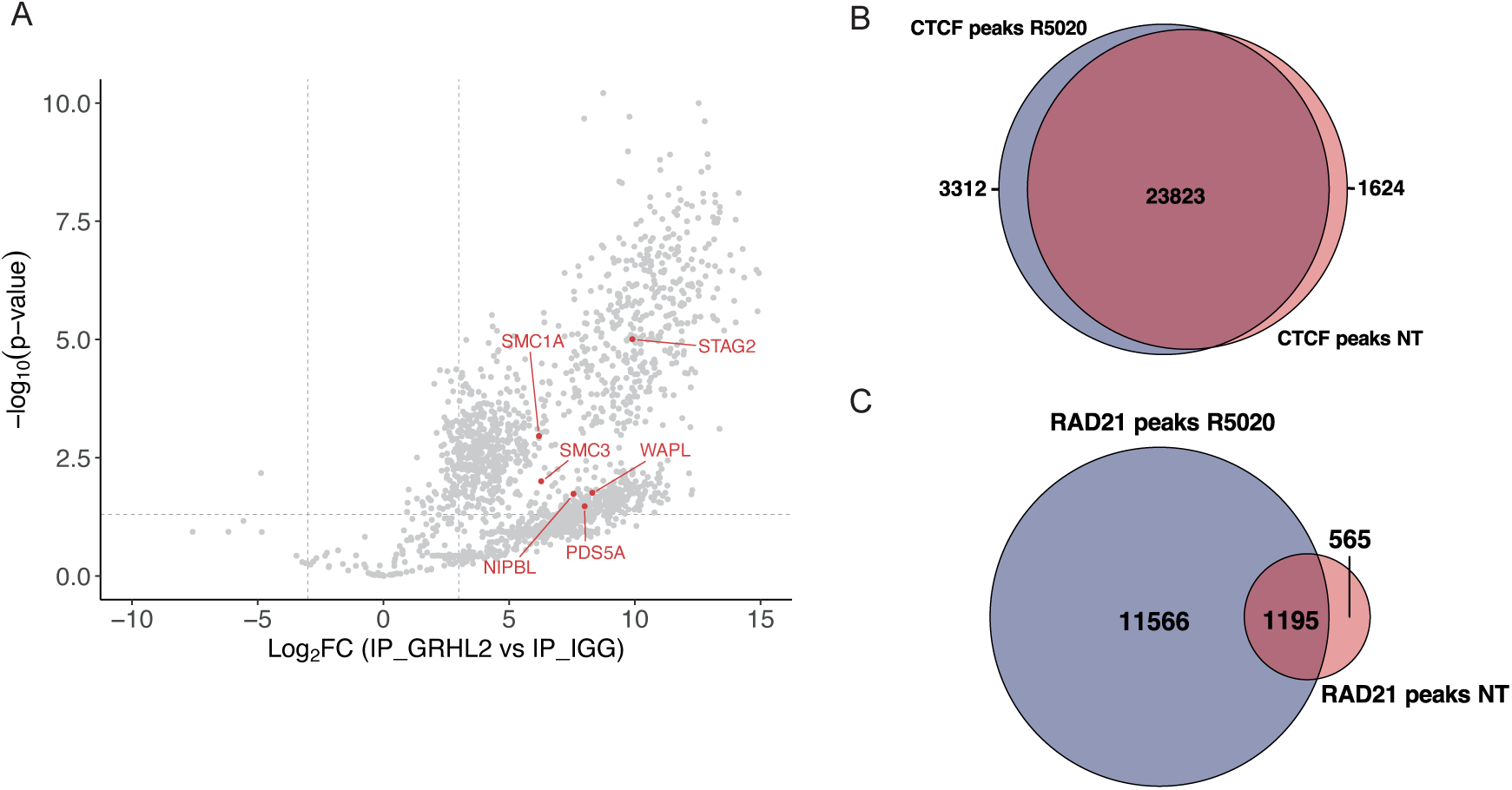
GRHL2 interacts with Cohesin complex members. A) Volcano plot depicting the results of the GRHL2 RIME of hormone depleted conditions vs. the IgG control. Each grey dot represents a single protein identified in mass spectrometry. Grey dotted lines represent Log2FC (−3,3) and -Log10(p-value) cutoffs (0.05) 1,140 proteins passed these criteria and were identified as a GRHL2 interactor. Highlighted dots in red are known Cohesin subunits, loading and releasing factors that are significantly interacting with GRHL2. B-C) Venn diagrams showing the overlap between non-treated and 30 minutes R5020 treated CTCF ChIP-seq peaks^49^ (B) or between non-treated and 30 minutes R5020 treated RAD21 ChIP-seq peaks^49^ (C) in T47D cells. Reanalysis of data (i.e. called peaks) from ChIP-Atlas. Peaks were considered overlapping if they overlapped by at least 1 bp.

## References

1. Klemm, S.L., Shipony, Z., and Greenleaf, W.J. (2019). Chromatin accessibility and the regulatory epigenome. Nat Rev Genet 20, 207–220. 10.1038/s41576-018-0089-8.

2. Ravasi, T., Suzuki, H., Cannistraci, C.V., Katayama, S., Bajic, V.B., Tan, K., Akalin, A., Schmeier, S., Kanamori-Katayama, M., Bertin, N., et al. (2010). An Atlas of Combinatorial Transcriptional Regulation in Mouse and Man. Cell 140, 744–752. 10.1016/j.cell.2010.01.044.

3. Fedorova, E., and Zink, D. (2008). Nuclear architecture and gene regulation. Biochim Biophys Acta 1783, 2174–2184. 10.1016/j.bbamcr.2008.07.018.

4. Bhardwaj, N., Yan, K.-K., and Gerstein, M.B. (2010). Analysis of diverse regulatory networks in a hierarchical context shows consistent tendencies for collaboration in the middle levels. Proceedings of the National Academy of Sciences 107, 6841– 6846. 10.1073/pnas.0910867107.

5. Kim, J., Choi, M., Kim, J.-R., Jin, H., Kim, V.N., and Cho, K.-H. (2012). The co-regulation mechanism of transcription factors in the human gene regulatory network. Nucleic Acids Res 40, 8849–8861. 10.1093/nar/gks664.

6. Nordin, A., Zambanini, G., Enar Jonasson, M., Weiss, T., van de Grift, Y., Pagella, P., and Cantù, C. (2025). Construction of an atlas of transcription factor binding during mouse development identifies popular regulatory regions. Development 152, dev204311. 10.1242/dev.204311.

7. The steroid receptor superfamily: mechanisms of diversity - Fuller - 1991 - The FASEB Journal - Wiley Online Library https://faseb.onlinelibrary.wiley.com/doi/10.1096/fasebj.5.15.1743440.

8. Frigo, D.E., Bondesson, M., and Williams, C. (2021). Nuclear receptors: from molecular mechanisms to therapeutics. Essays Biochem 65, 847–856. 10.1042/EBC20210020.

9. Brisken, C., Park, S., Vass, T., Lydon, J.P., O’Malley, B.W., and Weinberg, R.A. (1998). A paracrine role for the epithelial progesterone receptor in mammary gland development. Proc Natl Acad Sci U S A 95, 5076–5081.

10. Brisken, C., and Scabia, V. (2020). 90 YEARS OF PROGESTERONE: Progesterone receptor signaling in the normal breast and its implications for cancer. J Mol Endocrinol 65, T81–T94. 10.1530/JME-20-0091.

11. Ataca, D., Aouad, P., Constantin, C., Laszlo, C., Beleut, M., Shamseddin, M., Rajaram, R.D., Jeitziner, R., Mead, T.J., Caikovski, M., et al. (2020). The secreted protease Adamts18 links hormone action to activation of the mammary stem cell niche. Nat Commun 11, 1571. 10.1038/s41467-020-15357-y.

12. Bartlett, T.E., Evans, I., Jones, A., Barrett, J.E., Haran, S., Reisel, D., Papaikonomou, K., Jones, L., Herzog, C., Pashayan, N., et al. (2022). Antiprogestins reduce epigenetic field cancerization in breast tissue of young healthy women. Genome Medicine 14, 64. 10.1186/s13073-022-01063-5.

13. Zaret, K.S., and Carroll, J.S. (2011). Pioneer transcription factors: establishing competence for gene expression. Genes Dev. 25, 2227–2241. 10.1101/gad.176826.111.

14. Jozwik, K.M., and Carroll, J.S. (2012). Pioneer factors in hormone-dependent cancers. Nat Rev Cancer 12, 381–385. 10.1038/nrc3263.

15. Hurtado, A., Holmes, K.A., Ross-Innes, C.S., Schmidt, D., and Carroll, J.S. (2011). FOXA1 is a key determinant of estrogen receptor function and endocrine response. Nat Genet 43, 27–33. 10.1038/ng.730.

16. Carroll, J.S., Liu, X.S., Brodsky, A.S., Li, W., Meyer, C.A., Szary, A.J., Eeckhoute, J., Shao, W., Hestermann, E.V., Geistlinger, T.R., et al. (2005). Chromosome-wide mapping of estrogen receptor binding reveals long-range regulation requiring the forkhead protein FoxA1. Cell 122, 33–43. 10.1016/j.cell.2005.05.008.

17. Mohammed, H., D’Santos, C., Serandour, A.A., Ali, H.R., Brown, G.D., Atkins, A., Rueda, O.M., Holmes, K.A., Theodorou, V., Robinson, J.L.L., et al. (2013). Endogenous purification reveals GREB1 as a key estrogen receptor regulatory factor. Cell Rep 3, 342–349. 10.1016/j.celrep.2013.01.010.

18. Jozwik, K.M., Chernukhin, I., Serandour, A.A., Nagarajan, S., and Carroll, J.S. (2016). FOXA1 Directs H3K4 Monomethylation at Enhancers via Recruitment of the Methyltransferase MLL3. Cell Reports 17, 2715–2723. 10.1016/j.celrep.2016.11.028.

19. Eeckhoute, J., Keeton, E.K., Lupien, M., Krum, S.A., Carroll, J.S., and Brown, M. (2007). Positive Cross-Regulatory Loop Ties GATA-3 to Estrogen Receptor α Expression in Breast Cancer. Cancer Research 67, 6477–6483. 10.1158/0008-5472.CAN-07-0746.

20. Martin, E.M., Orlando, K.A., Yokobori, K., and Wade, P.A. (2021). The estrogen receptor/GATA3/FOXA1 transcriptional network: lessons learned from breast cancer. Curr Opin Struct Biol 71, 65–70. 10.1016/j.sbi.2021.05.015.

21. Reese, R.M., Harrison, M.M., and Alarid, E.T. (2019). Grainyhead-like Protein 2: The Emerging Role in Hormone-Dependent Cancers and Epigenetics. Endocrinology 160, 1275–1288. 10.1210/en.2019-00213.

22. Zheng, C., Allen, K.O., Liu, T., Solodin, N.M., Meyer, M.B., Salem, K., Tsourkas, P.K., McIlwain, S.J., Vera, J.M., Cromwell, E.R., et al. (2024). Elevated GRHL2 Imparts Plasticity in ER-Positive Breast Cancer Cells. Cancers 16, 2906. 10.3390/cancers16162906.

23. Chi, D., Singhal, H., Li, L., Xiao, T., Liu, W., Pun, M., Jeselsohn, R., He, H., Lim, E., Vadhi, R., et al. (2019). Estrogen receptor signaling is reprogrammed during breast tumorigenesis. Proc Natl Acad Sci U S A 116, 11437–11443. 10.1073/pnas.1819155116.

24. Wang, Z., Coban, B., Wu, H., Chouaref, J., Daxinger, L., Paulsen, M.T., Ljungman, M., Smid, M., Martens, J.W.M., and Danen, E.H.J. (2023). GRHL2-controlled gene expression networks in luminal breast cancer. Cell Commun Signal 21, 15. 10.1186/s12964-022-01029-5.

25. Reese, R.M., Helzer, K.T., Allen, K.O., Zheng, C., Solodin, N., and Alarid, E.T. (2022). GRHL2 Enhances Phosphorylated Estrogen Receptor (ER) Chromatin Binding and Regulates ER-Mediated Transcriptional Activation and Repression. Molecular and Cellular Biology 42, e00191–22. 10.1128/mcb.00191-22.

26. Cocce, K.J., Jasper, J.S., Desautels, T.K., Everett, L., Wardell, S., Westerling, T., Baldi, R., Wright, T.M., Tavares, K., Yllanes, A., et al. (2019). The Lineage Determining Factor GRHL2 Collaborates with FOXA1 to Establish a Targetable Pathway in Endocrine Therapy-Resistant Breast Cancer. Cell Reports 29, 889–903.e10. 10.1016/j.celrep.2019.09.032.

27. Wilanowski, T., Tuckfield, A., Cerruti, L., O’Connell, S., Saint, R., Parekh, V., Tao, J., Cunningham, J.M., and Jane, S.M. (2002). A highly conserved novel family of mammalian developmental transcription factors related to Drosophila grainyhead. Mech Dev 114, 37–50. 10.1016/s0925-4773(02)00046-1.

28. Jacobs, J., Atkins, M., Davie, K., Imrichova, H., Romanelli, L., Christiaens, V., Hulselmans, G., Potier, D., Wouters, J., Taskiran, I.I., et al. (2018). The transcription factor Grainy head primes epithelial enhancers for spatiotemporal activation by displacing nucleosomes. Nat Genet 50, 1011–1020. 10.1038/s41588-018-0140-x.

29. Ming, Q., Roske, Y., Schuetz, A., Walentin, K., Ibraimi, I., Schmidt-Ott, K.M., and Heinemann, U. (2018). Structural basis of gene regulation by the Grainyhead/CP2 transcription factor family. Nucleic Acids Res 46, 2082–2095. 10.1093/nar/gkx1299.

30. Venkatesan, K., McManus, H.R., Mello, C.C., Smith, T.F., and Hansen, U. (2003). Functional conservation between members of an ancient duplicated transcription factor family, LSF/Grainyhead. Nucleic Acids Res 31, 4304–4316. 10.1093/nar/gkg644.

31. Mohammed, H., Taylor, C., Brown, G.D., Papachristou, E.K., Carroll, J.S., and D’Santos, C.S. (2016). Rapid immunoprecipitation mass spectrometry of endogenous proteins (RIME) for analysis of chromatin complexes. Nat Protoc 11, 316–326. 10.1038/nprot.2016.020.

32. MacFawn, I., Wilson, H., Selth, L.A., Leighton, I., Serebriiskii, I., Bleackley, R.C., Elzamzamy, O., Farris, J., Pifer, P.M., Richer, J., et al. (2019). Grainyhead-like-2 confers NK-sensitivity through interactions with epigenetic modifiers. Molecular Immunology 105, 137–149. 10.1016/j.molimm.2018.11.006.

33. Ramamurthy, L., Barbour, V., Tuckfield, A., Clouston, D.R., Topham, D., Cunningham, J.M., and Jane, S.M. (2001). Targeted disruption of the CP2 gene, a member of the NTF family of transcription factors. J Biol Chem 276, 7836–7842. 10.1074/jbc.M004351200.

34. Thomas, P.D., Ebert, D., Muruganujan, A., Mushayahama, T., Albou, L.-P., and Mi, H. (2022). PANTHER: Making genome-scale phylogenetics accessible to all. Protein Sci 31, 8–22. 10.1002/pro.4218.

35. Mohammed, H., Russell, I.A., Stark, R., Rueda, O.M., Hickey, T.E., Tarulli, G.A., Serandour, A.A., Birrell, S.N., Bruna, A., Saadi, A., et al. (2015). Progesterone receptor modulates ERα action in breast cancer. Nature 523, 313–317. 10.1038/nature14583.

36. Zaurin, R., Ferrari, R., Nacht, A.S., Carbonell, J., Le Dily, F., Font-Mateu, J., de Llobet Cucalon, L.I., Vidal, E., Lioutas, A., Beato, M., et al. (2021). A set of accessible enhancers enables the initial response of breast cancer cells to physiological progestin concentrations. Nucleic Acids Res 49, 12716–12731. 10.1093/nar/gkab1125.

37. Robinson, J.T., Thorvaldsdóttir, H., Winckler, W., Guttman, M., Lander, E.S., Getz, G., and Mesirov, J.P. (2011). Integrative genomics viewer. Nat Biotechnol 29, 24–26. 10.1038/nbt.1754.

38. Wang, Q., Li, M., Wu, T., Zhan, L., Li, L., Chen, M., Xie, W., Xie, Z., Hu, E., Xu, S., et al. (2022). Exploring Epigenomic Datasets by ChIPseeker. Curr Protoc 2, e585. 10.1002/cpz1.585.

39. Yu, G., Wang, L.-G., and He, Q.-Y. (2015). ChIPseeker: an R/Bioconductor package for ChIP peak annotation, comparison and visualization. Bioinformatics 31, 2382–2383. 10.1093/bioinformatics/btv145.

40. Heinz, S., Benner, C., Spann, N., Bertolino, E., Lin, Y.C., Laslo, P., Cheng, J.X., Murre, C., Singh, H., and Glass, C.K. (2010). Simple combinations of lineage-determining transcription factors prime cis-regulatory elements required for macrophage and B cell identities. Mol Cell 38, 576–589. 10.1016/j.molcel.2010.05.004.

41. Xu, S., Hu, E., Cai, Y., Xie, Z., Luo, X., Zhan, L., Tang, W., Wang, Q., Liu, B., Wang, R., et al. (2024). Using clusterProfiler to characterize multiomics data. Nat Protoc 19, 3292–3320. 10.1038/s41596-024-01020-z.

42. Yao, L., Berman, B.P., and Farnham, P.J. (2015). Demystifying the secret mission of enhancers: linking distal regulatory elements to target genes. Crit Rev Biochem Mol Biol 50, 550–573. 10.3109/10409238.2015.1087961.

43. Sanyal, A., Lajoie, B.R., Jain, G., and Dekker, J. (2012). The long-range interaction landscape of gene promoters. Nature 489, 109–113. 10.1038/nature11279.

44. Li, G., Ruan, X., Auerbach, R.K., Sandhu, K.S., Zheng, M., Wang, P., Poh, H.M., Goh, Y., Lim, J., Zhang, J., et al. (2012). Extensive promoter-centered chromatin interactions provide a topological basis for transcription regulation. Cell 148, 84–98. 10.1016/j.cell.2011.12.014.

45. Mifsud, B., Tavares-Cadete, F., Young, A.N., Sugar, R., Schoenfelder, S., Ferreira, L., Wingett, S.W., Andrews, S., Grey, W., Ewels, P.A., et al. (2015). Mapping long-range promoter contacts in human cells with high-resolution capture Hi-C. Nat Genet 47, 598–606. 10.1038/ng.3286.

46. Tettey, T.T., Rinaldi, L., and Hager, G.L. (2023). Long-range gene regulation in hormone-dependent cancer. Nat Rev Cancer 23, 657–672. 10.1038/s41568-023-00603-4.

47. Jin, F., Li, Y., Dixon, J.R., Selvaraj, S., Ye, Z., Lee, A.Y., Yen, C.-A., Schmitt, A.D., Espinoza, C.A., and Ren, B. (2013). A high-resolution map of the three-dimensional chromatin interactome in human cells. Nature 503, 290–294. 10.1038/nature12644.

48. Handoko, L., Xu, H., Li, G., Ngan, C.Y., Chew, E., Schnapp, M., Lee, C.W.H., Ye, C., Ping, J.L.H., Mulawadi, F., et al. (2011). CTCF-mediated functional chromatin interactome in pluripotent cells. Nat Genet 43, 630–638. 10.1038/ng.857.

49. Ramírez-Cuéllar, J., Ferrari, R., Sanz, R.T., Valverde-Santiago, M., García-García, J., Nacht, A.S., Castillo, D., Le Dily, F., Neguembor, M.V., Malatesta, M., et al. (2024). LATS1 controls CTCF chromatin occupancy and hormonal response of 3D-grown breast cancer cells. EMBO J 43, 1770–1798. 10.1038/s44318-024-00080-x.

50 Le Dily, F., Baù, D., Pohl, A., Vicent, G.P., Serra, F., Soronellas, D., Castellano, G., Wright, R.H.G., Ballare, C., Filion, G., et al. (2014). Distinct structural transitions of chromatin topological domains correlate with coordinated hormone-induced gene regulation. Genes Dev 28, 2151–2162. 10.1101/gad.241422.114.

51. Holding, A.N., Giorgi, F.M., Donnelly, A., Cullen, A.E., Nagarajan, S., Selth, L.A., and Markowetz, F. (2019). VULCAN integrates ChIP-seq with patient-derived co-expression networks to identify GRHL2 as a key co-regulator of ERa at enhancers in breast cancer. Genome Biol 20, 91. 10.1186/s13059-019-1698-z.

52. Khaled, W.T., Read, E.K.C., Nicholson, S.E., Baxter, F.O., Brennan, A.J., Came, P.J., Sprigg, N., McKenzie, A.N.J., and Watson, C.J. (2007). The IL-4/IL-13/Stat6 signalling pathway promotes luminal mammary epithelial cell development. Development 134, 2739–2750. 10.1242/dev.003194.

53. Oueslati, M., Bettaieb, I., Ben Younes, R., Gamoudi, A., Rahal, K., and Oueslati, R. (2022). STAT-5 and STAT-6 in Breast Cancer: Potential Crosstalk With Estrogen and Progesterone Receptors Can Affect Cell Proliferation and Metastasis. J Clin Med Res 14, 416–424. 10.14740/jocmr4785.

54. Abu-Farha, M., Lambert, J.-P., Al-Madhoun, A.S., Elisma, F., Skerjanc, I.S., and Figeys, D. (2008). The tale of two domains: proteomics and genomics analysis of SMYD2, a new histone methyltransferase. Mol Cell Proteomics 7, 560–572. 10.1074/mcp.M700271-MCP200.

55. Zhang, X., Tanaka, K., Yan, J., Li, J., Peng, D., Jiang, Y., Yang, Z., Barton, M.C., Wen, H., and Shi, X. (2013). Regulation of estrogen receptor α by histone methyltransferase SMYD2-mediated protein methylation. Proc Natl Acad Sci U S A 110, 17284–17289. 10.1073/pnas.1307959110.

56. Yang, L., Kumegawa, K., Saeki, S., Nakadai, T., and Maruyama, R. (2024). Identification of lineage-specific epigenetic regulators FOXA1 and GRHL2 through chromatin accessibility profiling in breast cancer cell lines. Cancer Gene Ther, 1–10. 10.1038/s41417-024-00745-z.

57. Bernardo, G.M., Bebek, G., Ginther, C.L., Sizemore, S.T., Lozada, K.L., Miedler, J.D., Anderson, L.A., Godwin, A.K., Abdul-Karim, F.W., Slamon, D.J., et al. (2013). FOXA1 represses the molecular phenotype of basal breast cancer cells. Oncogene 32, 554–563. 10.1038/onc.2012.62.

58. Bernardo, G.M., Lozada, K.L., Miedler, J.D., Harburg, G., Hewitt, S.C., Mosley, J.D., Godwin, A.K., Korach, K.S., Visvader, J.E., Kaestner, K.H., et al. (2010). FOXA1 is an essential determinant of ERα expression and mammary ductal morphogenesis. Development 137, 2045–2054. 10.1242/dev.043299.

59. Liu, Y., Zhao, Y., Skerry, B., Wang, X., Colin-Cassin, C., Radisky, D.C., Kaestner, K.H., and Li, Z. (2016). Foxa1 is essential for mammary duct formation. genesis 54, 277–285. 10.1002/dvg.22929.

60. Ceballos-Chávez, M., Subtil-Rodríguez, A., Giannopoulou, E.G., Soronellas, D., Vázquez-Chávez, E., Vicent, G.P., Elemento, O., Beato, M., and Reyes, J.C. (2015). The chromatin Remodeler CHD8 is required for activation of progesterone receptor-dependent enhancers. PLoS Genet 11, e1005174. 10.1371/journal.pgen.1005174.

61. van de Grift, Y.B., Aarts, M.T., Wiese, K.E., Heijmans, N., Hooijkaas, I.B., Pritchard, C.E., Henneman, L., Krimpenfort, P.J., van Boxtel, A.L., and van Amerongen, R. (2025). GRHL and PGR control WNT4 expression in the mammary gland via 3D looping of conserved and species-specific enhancers. bioRxiv, 2025.04.11.648333. 10.1101/2025.04.11.648333.

62. Hughes, C.S., Foehr, S., Garfield, D.A., Furlong, E.E., Steinmetz, L.M., and Krijgsveld, J. (2014). Ultrasensitive proteome analysis using paramagnetic bead technology. Mol Syst Biol 10, 757. 10.15252/msb.20145625.

63 Aarts, M.T., Wagner, M., van der Wal, T., van Boxtel, A.L., and van Amerongen, R. (2023). A molecular toolbox to study progesterone receptor signaling. J Mammary Gland Biol Neoplasia 28, 24. 10.1007/s10911-023-09550-0.

64. Ewels, P., Magnusson, M., Lundin, S., and Käller, M. (2016). MultiQC: summarize analysis results for multiple tools and samples in a single report. Bioinformatics 32, 3047–3048. 10.1093/bioinformatics/btw354.

65. 65. Trimmomatic: a flexible trimmer for Illumina sequence data | Bioinformatics | Oxford Academic https://academic.oup.com/bioinformatics/article/30/15/2114/2390096.

66. Kim, D., Paggi, J.M., Park, C., Bennett, C., and Salzberg, S.L. (2019). Graph-based genome alignment and genotyping with HISAT2 and HISAT-genotype. Nat Biotechnol 37, 907–915. 10.1038/s41587-019-0201-4.

67. Anders, S., Pyl, P.T., and Huber, W. (2015). HTSeq--a Python framework to work with high-throughput sequencing data. Bioinformatics 31, 166–169. 10.1093/bioinformatics/btu638.

68. Love, M.I., Huber, W., and Anders, S. (2014). Moderated estimation of fold change and dispersion for RNA-seq data with DESeq2. Genome Biology 15, 550. 10.1186/s13059-014-0550-8.

69. Chua, S.L., See Too, W.C., Khoo, B.Y., and Few, L.L. (2011). UBC and YWHAZ as suitable reference genes for accurate normalisation of gene expression using MCF7, HCT116 and HepG2 cell lines. Cytotechnology 63, 645–654. 10.1007/s10616-011-9383-4.

70. Zambanini, G., Nordin, A., Jonasson, M., Pagella, P., and Cantù, C. (2022). A new CUT&RUN low volume-urea (LoV-U) protocol optimized for transcriptional co-factors uncovers Wnt/β-catenin tissue-specific genomic targets. Development 149, dev201124. 10.1242/dev.201124.

71. de Sena Brandine, G., and Smith, A.D., (2019). Falco: high-speed FastQC emulation for quality control of sequencing data. F1000Res 8, 1874. 10.12688/f1000research.21142.2.

72. Bushnell, B., Rood, J., and Singer, E. (2017). BBMerge – Accurate paired shotgun read merging via overlap. PLOS ONE 12, e0185056. 10.1371/journal.pone.0185056.

73. Langmead, B., and Salzberg, S.L. (2012). Fast gapped-read alignment with Bowtie 2. Nat Methods 9, 357–359. 10.1038/nmeth.1923.

74. Li, H., Handsaker, B., Wysoker, A., Fennell, T., Ruan, J., Homer, N., Marth, G., Abecasis, G., and Durbin, R. (2009). The Sequence Alignment/Map format and SAMtools. Bioinformatics 25, 2078–2079. 10.1093/bioinformatics/btp352.

75. Quinlan, A.R., and Hall, I.M. (2010). BEDTools: a flexible suite of utilities for comparing genomic features. Bioinformatics 26, 841–842. 10.1093/bioinformatics/btq033.

76. Nordin, A., Zambanini, G., Pagella, P., and Cantù, C. (2023). The CUT&RUN suspect list of problematic regions of the genome. Genome Biology 24, 185. 10.1186/s13059-023-03027-3.

77. Zhang, Y., Liu, T., Meyer, C.A., Eeckhoute, J., Johnson, D.S., Bernstein, B.E., Nusbaum, C., Myers, R.M., Brown, M., Li, W., et al. (2008). Model-based Analysis of ChIP-Seq (MACS). Genome Biology 9, R137. 10.1186/gb-2008-9-9-r137.

78. Bolger, A.M., Lohse, M., and Usadel, B. (2014). Trimmomatic: a flexible trimmer for Illumina sequence data. Bioinformatics 30, 2114–2120. 10.1093/bioinformatics/btu170.

79. Amemiya, H.M., Kundaje, A., and Boyle, A.P. (2019). The ENCODE Blacklist: Identification of Problematic Regions of the Genome. Sci Rep 9, 9354. 10.1038/s41598-019-45839-z.

80. Ramírez, F., Dündar, F., Diehl, S., Grüning, B.A., and Manke, T. (2014). deepTools: a flexible platform for exploring deep-sequencing data. Nucleic Acids Res 42, W187–191. 10.1093/nar/gku365.

81. Zou, Z., Ohta, T., and Oki, S. (2024). ChIP-Atlas 3.0: a data-mining suite to explore chromosome architecture together with large-scale regulome data. Nucleic Acids Res 52, W45–W53. 10.1093/nar/gkae358.

